# Spatiotemporal Genetic Diversity of Lions

**DOI:** 10.1101/2020.01.07.896431

**Authors:** Caitlin J. Curry, Brian W. Davis, Laura D. Bertola, Paula A. White, William J. Murphy, James N. Derr

## Abstract

The Scramble for Africa in the late 1800s marked the beginning of increased human population growth in Africa. Here, we determined the genetic architecture of both historical and modern lions to identify changes in genetic diversity that occurred during this period of landscape and anthropogenic change. We surveyed microsatellite and mitochondrial genetic variation from 143 high-quality museum specimens of known provenance and combined them with data from recently published nuclear and mitochondrial studies. Analysis of variation at 9 microsatellites and 280 polymorphic mitogenome SNPs indicate the presence of male-mediated gene flow and recent isolation of local subpopulations, likely due to habitat fragmentation. Nuclear markers showed a significant decrease in genetic diversity from the historical (H_E_=0.833) to the modern (H_E_=0.796) populations, while mitochondrial genetic diversity was maintained (Hd=0.98 for both). While the historical population appears to have been panmictic based on nDNA data, hierarchical structure analysis identified four tiers of fine structure in modern populations, able to detect most sampling locations. Mitochondrial analyses identified 4 clusters: Southern, Mixed, Eastern, and Western; and were consistent between modern and historically sampled haplotypes. Within the last century, habitat fragmentation caused lion subpopulations to become more isolated as human expansion changed the African landscape. This resulted in an increase in fine-scale nuclear genetic structure and loss of genetic diversity as subpopulations became more differentiated, while mitochondrial structure and diversity was maintained over time.

## INTRODUCTION

The Scramble for Africa in the late-1800’s increased European control of the African continent from 10% to 90% (1). The influence resultant of European settlement inevitably lead to an exponential increase in the human population, urban development, and rural expansion (2) resulting in changes to the African landscape and fragmentation to once continuous wildlife ranges (3–5).

Multiple published investigations document the genetic consequences of large scale landscape changes over short periods of time (6–9) (i.e. 100 years). Levels of genetic diversity are directly related to a species ability to adapt, survive, and thrive; and loss of genetic diversity can be detrimental to overall population health and long-term survival (10–13). Many species across Africa are declining due to human-induced threats (14–16), even in protected areas (16). The adverse effects of low genetic diversity have been observed in small feline populations that exist in heavily managed fenced reserves (17–21). Historically, the lion (*Panthera leo*) range was more continuous and connected (22). The lion population has changed dramatically over the past 100 years (3, 22, 23), particularly in terms of population size and distribution in response to habitat availability and anthropogenic pressures related to a growing human population (24–27).

Around the turn of the 19th century, explorers, naturalists and hunters went on expeditions to collect biological specimens for preservation in natural history museums. These expeditions resulted in hundreds of lion specimens being deposited in museums across the world that predate the precipitous human population growth across Africa (1, 2). With the continued development of techniques for improved isolation and sequencing of degraded genetic material (ancient DNA, aDNA), these collections now provide access to genetic information from historically sampled individuals as well as their contemporary counterparts.

Previous studies that sampled nuclear genetic diversity reported both high (28–30) and low (31–33) levels of gene flow, but this was largely dependent on the amount of connectivity present between sampling locations. For example, isolated populations such as those in the Kainji Lake National Park from Yankari Game Reserve in Nigeria (31) and Kafue National Park from the Luangwa Valley Ecosystem in Zambia (32). Genetic differentiation can even be seen between populations within national parks (33). However, where there are no geographic or man-made barriers to limit movement, there is only weak evidence of population structure and high levels of gene flow (28, 30).

Studies including historical and ancient lion samples have been primarily restricted to mtDNA analyses incorporated within a modern lion dataset (22, 34–37). A recent study including historical individuals focused on assessing changes in the recent past but was confined to a local analysis of the Kavango–Zambezi transfrontier conservation area (KAZA) (38). Here, we report the first range-wide study assessing changes in genetic diversity of the lion, based on both historical and modern samples collected throughout African and India. By comparing diversity estimates from samples from different time periods we can detect and evaluate changes in genetic diversity that have occurred during this time of landscape and anthropogenic change.

## RESULTS

### Nuclear DNA

The modern dataset (MD) contained 135 lions from 14 sampling locations and the historical dataset (HD) consisted of 143 lions (SI Appendix S1). Nine microsatellite loci (Leo006, Leo008, Leo085, Leo098, Leo126, Leo224, Leo230, Leo247, Leo281) were shared between with two datasets. The MD had greater than 75% allele calls reported, and the HD had an average of 90% amplification success across the nine loci. Sampling was similarly distributed across the lion range for both the MD and HD (Figure 1).

**Figure 1.**
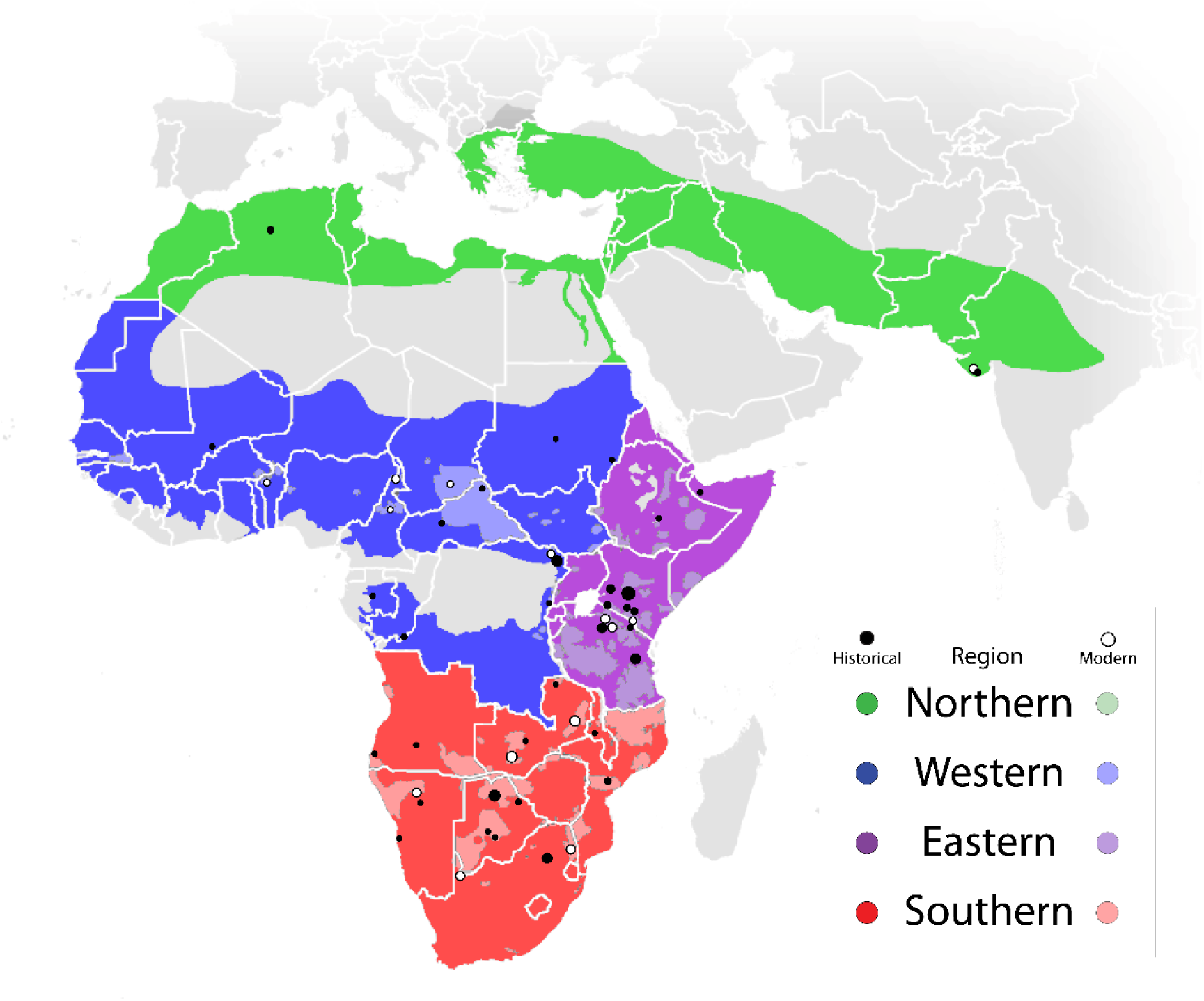
Map of lion sample locations. Dot size coincides with sample size for each location.

Fine-scale structure was observed in the MD but not the HD. The MD hierarchical structure analysis resulted in four tiers of structure: Continental (K=2), Subcontinental (K=4), Regional (K=6), and Local (K=11), as seen in the final analysis with location priors (Figure 2, SI Appendix S2 shows a graphical display of the step-by-step hierarchical population structure). The initial run had a ΔK of 2 separating Asia (GIR) from Africa. The African population could then be further broken down into a Western, Eastern, and Southern population with a ΔK of 3 from structural analysis of only the African population. Analysis of the Western population also resulted a ΔK of 3 separating a West African (WES), Central African (CEN) and a population between the two (MID). The Eastern population had a ΔK of 2 separating lions in Kenya (KEN) from all lions sampled in Tanzania (TAN). The Southern population is separated into 5 local populations that can be grouped into 3 regional populations as ΔK was 5, however, there was a sizable peak also seen at K=3. Eastern and western Zambia (ZAE and ZAW) make up a Southeast population while Etosha and Kalahari (ETO & KAL) make up a Southwest population. Kruger was identified as a single population at both K=3 and K=5. Population clustering in the MD principal coordinate analysis (PCoA) follows the Subcontinental tier (Figure 3). Mean heterozygosity across polymorphic loci (H_E_) is lowest in GIR and highest in CEN although only 44% of loci are polymorphic in this population (SI Appendix S3).

**Figure 2.**
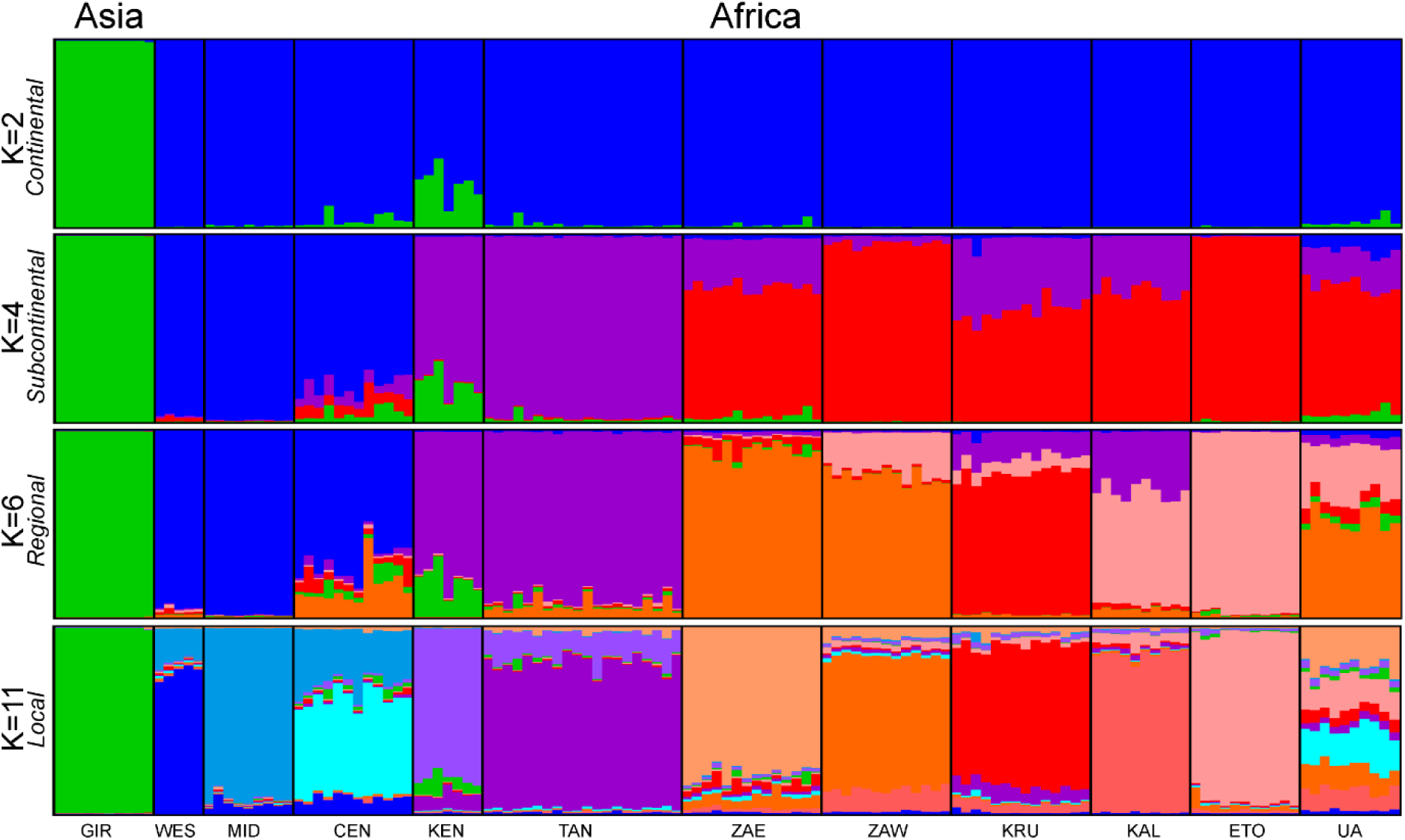
The four tiers of lion population structure as determined by hierarchical structure analysis based on 9 microsatellites. Groups are colored based on Figure 1.

**Figure 3.**
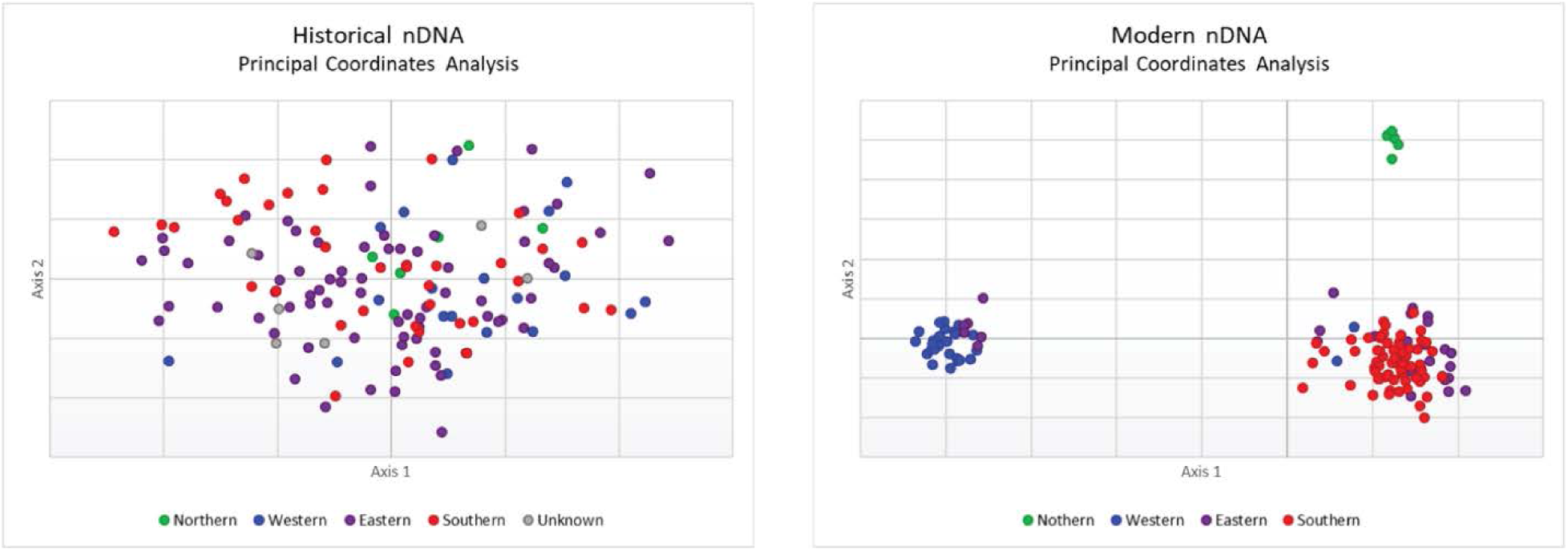
Results of a principal coordinate analysis (PCoA) of 9 microsatellite loci analyzed in historical and contemporary lion samples.

Bayesian clustering in STRUCTURE did not identify any population structure in the HD. While ΔK was also 2 for the initial run, individuals could not be assigned to meaningful populations. Further evidence of this lack of structure was observed in the PCoA results (Figure 3). PCoA did not reveal any population clusters but did show weak evidence of isolation-by-distance (IBD) (SI Appendix S5), indicating an admixed population.

A significant decrease (p-value < 0.005) from HD to MD was evident across diversity indices (Table 1). Correcting for sample size through rarefaction, the HD has an allelic richness of 14.2 and private allelic richness of 4.6, higher than the MD at 11.3 and 1.7, respectively. The Garza-Williamson index (M) of the HD is 0.41 and the MD is 0.32.

**Table 1.** Historical versus modern genetic diversity for nDNA and mtDNA. N = Sample Size, HE = Expected Heterozygosity, A = Allelic Richness, PA = Private Alleles, M = Garza-Williamson Index, s = Segregating Sites, π = Nucleotide Diversity, Hd = Haplotype Diversity, H = Number of Haplotypes, PM = Private Mutations. Trend is based on statistical significance from a comparison of means. HE, A, PA, and M had a p-value < 0.005 (***) indicating a downward trend (↓) from historical to modern. The p-value for π and Hd was > 0.05 (n.s.) indicating maintained diversity (→) from historical to modern.

|  | Historical | Modern | Sig. | Trend |
| --- | --- | --- | --- | --- |
| **nDNA**<br>(9 msat loci) | *N* 143 | *N* 135 |  |  |
|  | *H<sub>E</sub>* 0.833 | *H<sub>E</sub>* 0.796 | *** | ↓ |
|  | *A* 15 | *A* 11.6 | *** |  |
|  | *PA* 6.2 | *PA* 1.2 | *** |  |
|  | *M* 0.41 | *M* 0.32 | *** |  |
| **mtDNA**<br>(280 SNPs) | *N* 102 | *N* 19 |  |  |
|  | *s* 280 | *s* 258 |  |  |
|  | *π* 0.222 | *π* 0.258 | n.s. | → |
|  | *Hd* 0.98 | *Hd* 0.98 | n.s. |  |
|  | *H* 74 | *H* 17 |  |  |
|  | *PM* 22 | *PM* 1 |  |  |

### Mitochondrial DNA

Results from mitochondrial analyses can be found in Table 1. There was no significant difference found between mtDNA nucleotide diversity (π) and haplotype diversity (Hd). The historical population had 74 haplotypes with 22 private mutations while the modern population had 17 haplotypes with only 1 private mutation. Two haplotypes (Hap_33 and Hap_66) are in both datasets. The low number of private mutations (PM) in the modern population was likely a result of the small number of mitogenomes compared to the historical population. While we were able to obtain a large number of historical mitogenomes from museum samples, the number of modern mitogenomes was restricted to published data

Mitochondrial genome analyses identified 4 major clades: Southern, Mixed, Eastern and Western. Each clade was represented by at least one of the 19 modern lions. A Northern subclade was nested within the Western clade. Bootstrap values in the unrooted maximum likelihood (ML) tree (Figure 4) indicating strong support for these four clades. The 4 clusters in the PCA (SI Appendix S6) and the four main branches of the haplotype network (SI Appendix S7) also support these same four clades.

**Figure 4.**
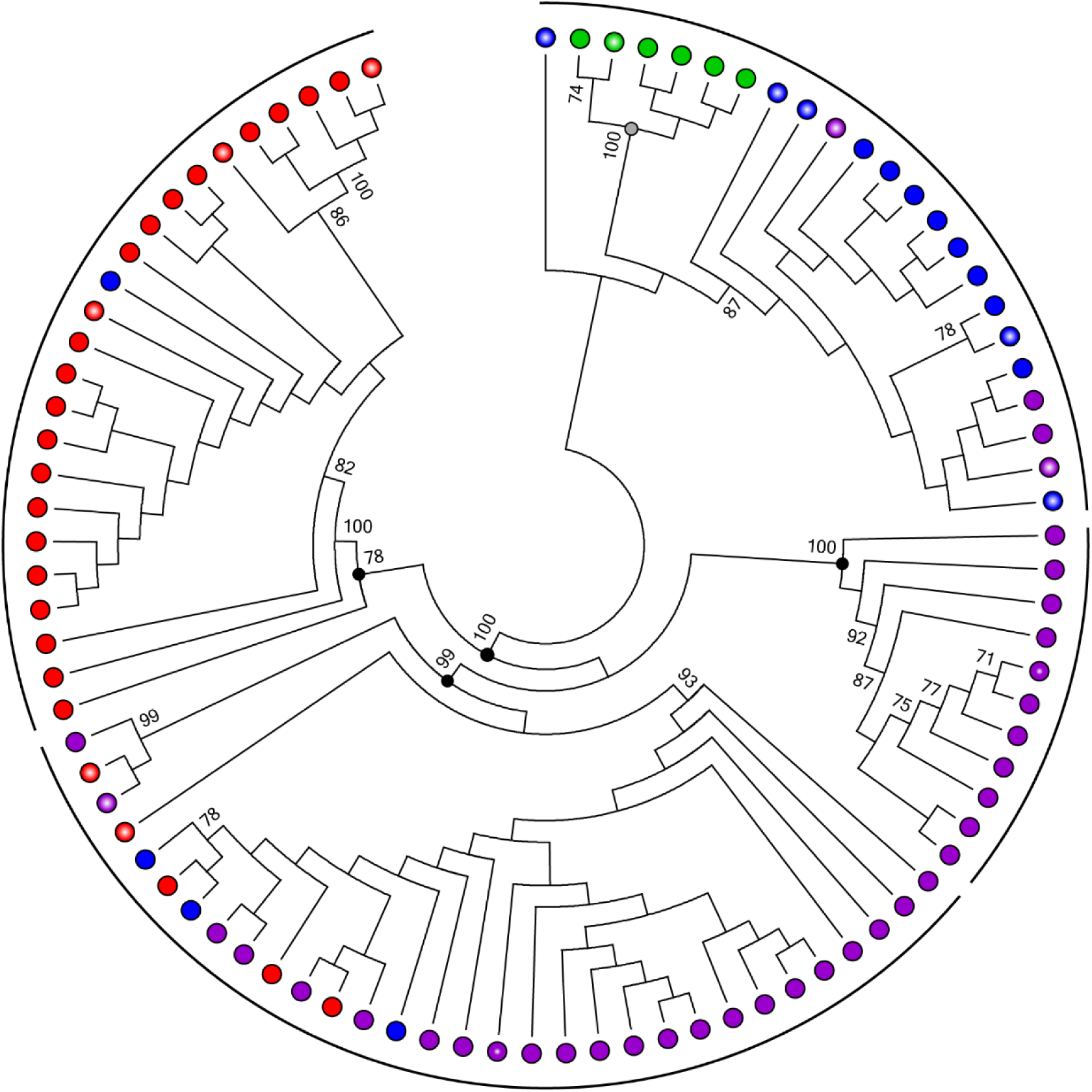
Maximum likelihood tree based on 280 variable sites in 121 lion mitogenomes showing nodes with >70% bootstrap support. Black dots denote the nodes of the four major clades. Arcs indicate clade boundaries. The hollow dot denotes the nested Gir Forest clade. Color corresponds to sampling location according to conventionally recognized regions in Figure 1.

There are five conventionally recognized regions of Africa according to the United Nations geoscheme for Africa: Southern, Eastern, Western, Central, and Northern (https://unstats.un.org/unsd/methodology/m49/). The Southern clade includes the conventionally recognized regions of Southern Africa incorporating Botswana, South Africa, Namibia and Zimbabwe. Haplotypes from Botswana and South Africa were present in both the Southern and Mixed clades. The Mixed clade consists of haplotypes from the Southern, Eastern, and Western subcontinental groups.

The Western clade included countries in the conventionally recognized regions of Central and Western Africa including present-day Democratic Republic of Congo, Benin, Central African Republic, and Cameroon. The Eastern clade consists primarily of historical lions from British East Africa, specifically present-day Kenya, Tanzania and Uganda, and a modern lion from Somalia. Eastern haplotypes within the Western clade are from bordering countries suggesting gene flow between neighboring regions, which is in line with previously published patterns (37).

The Mixed clade is intermediary, consisting of lions from Southern and Eastern Africa. Zambia and Malawi were found exclusively in the Mixed clade, while all other countries found in the Mixed clade are also within the Southern and/or Western clades. The historical samples from the Congo Region (present day Republic of the Congo and Gabon) were assigned to the Western subcontinental group by convention (Figure 1), but mitochondrial analyses consistently clustered them within the Southern and Mixed clades. The Congo Region is below the Congolese rainforests, a geographical barrier isolating them from West and Central Africa to the north. For the lion, the Congo Region is, therefore, closer to East and Southern Africa.

## DISCUSSION

While lions currently exhibit fine population structure, the historical population lacked any structure, suggesting lions acted as a panmictic population only a century ago. This result was supported by structure analysis and PCoA. The modern populations cluster into four subcontinental groups, which are not recovered in the historical population, even between Asia and Africa.

Four tiers of modern nuclear genetic structure were identified through hierarchical structure analysis. Continental structure separated Asia from Africa. The subcontinental tier identified three main groups in Africa: Western, Eastern, and Southern. Bertola *et al*. 2015 (39) also observed strong differentiation between Africa and Asia as well as subcontinental structure within Africa. Smitz *et al*. 2018 (30) identified only two groups at the subcontinental scale; lack of identification of a Southern group was likely due to low sampling. The regional tier divides the Southern group into a Southwest, South, and Southeast group. The highest level of population structure was able to detect most sampling locations, as in Antunes *et al*. 2008 (40). Only sampling locations in Tanzania were unable to be individually identified, similar to findings of Smitz *et al*. 2018 (30). Admixture was evident within local groups (Figure 2, K=11) indicating recent gene flow. The UA group is comprised of individuals sampled within local groups that could not be assigned to a particular tier due to admixture. Other range-wide studies of lions have shown a similar localized structure pattern with individuals assigned to sampling populations with evidence of isolation-by-distance (40, 41).

Habitat fragmentation restricts gene flow and often leads to loss of genetic diversity (42, 43). The Kavango–Zambezi transfrontier conservation area has seen a decrease in genetic and allelic diversity over the past century (38). Our study showed there has been a significant decrease in nDNA diversity from the range-wide historical to the modern lion population (Table 1). Expected heterozygosity, allelic richness, and number of private alleles have all significantly decreased (p-value < 0.0005). While there is evidence of a recent reduction in size in both the historical and modern populations both displaying an M value <0.67 (44) (Table 1), M is significantly lower in the modern population. This indicates the reduction predates the Scramble for Africa but has increased in the past century.

Given the strong signal differentiating the Asiatic and African lions in the analysis of modern populations, we predicted the historical lions from the Gir Forest National Park (NP) in India would cluster independently of the historical African lions. The PCoA, however, showed that the Gir Forest lions cluster together in the center of a single historical lion cluster (Figure 3). Lions were at the brink of extinction in Asia at the beginning of the 20th century (45) when these samples were collected (1906-1929). Today there are over 400 lions in the Gir Forest NP. The historical and modern samples from the Gir Forest NP were collected at the peak of a recent bottleneck and its subsequent population restoration. Our comparisons document this severe bottleneck that resulted in low genetic diversity in Asia compared to Africa (SI Appendix S3). Habitat fragmentation leading to the isolation of subpopulations within Africa appears to be following the same trend as the Asiatic lion a century ago.

Historical and modern mtDNA show strikingly different patterns than nuclear data. Although nuclear diversity has decreased significantly, mtDNA diversity has remained constant over time (Table 1). Mitochondrial DNA is matrilineally inherited, and localized studies show there is little or no female-mediated gene flow between subpopulations across Africa (32, 46, 47). Female lions primarily remain with their natal pride while males disperse (48, 49). Therefore, pride structure can dictate mtDNA population structure. With females remaining close to their natal prides, habitat fragmentation will not greatly alter pride structure, keeping mtDNA diversity constant over time. Accordingly, the distribution of modern haplotypes within clades is geographically consistent with historical haplotypes. The four major clades (Southern, Mixed, Eastern, and Western) geographically follow the subcontinental groups identified by nDNA analysis (Southern, Eastern, Western, and Northern).

The Western clade includes lions from West and Central Africa as well as Asiatic lions. Previous studies have suggested that the Asiatic and West and Central African lions should be grouped taxonomically (37, 50). United States Fish and Wildlife Services recently updated the lion taxonomy under the Endangered Species Act to recognize these populations as the same subspecies, *Panthera leo leo*, and East and Southern Africa populations as *Panthera leo melanochaita* (51). Our mitogenome results support this dichotomy, placing the Gir Forest NP lions within the Western clade in all analyses

## CONCLUSION

A century ago, the lion population consisted of close-proximity prides with enough overlapping movement to appear panmictic. Our comparison of nDNA and mtDNA analyses between historical and modern datasets indicates the presence of substantial historical male-mediated gene flow and evidence of recent isolation of local subpopulations due to habitat fragmentation. In the lion mating system, females primarily remain with their natal pride while males disperse (48, 49). If unobstructed by geographic or artificial barriers, a male home range can be hundreds to thousands of km2 (52–54) and can span different habitats (55–57). However, the original range of the lion has been severely reduced as a direct result of the growing human population (24) and changes in land-use, such as expansion of large-scale cultivation and increased movement of livestock into protected areas (58–60). As lion habitat has become more fragmented and groups of prides become more isolated, gene flow is restricted, and subpopulations become more genetically distinguishable. The dichotomy between the historic nuclear and mitochondrial structure is indicative of male-mediated gene flow and female philopatry. This pattern is not as evident in the modern population because fragmentation has hindered the ability of males to migrate between isolated subpopulations.

The differences evident between the historical and modern lion populations have several important conservation implications. If left unattended, these subpopulations could become completely isolated leading to further differentiation and reduction in genetic diversity. Managing a species as a continuous population without a continuous habitat requires considerable resources. Lions currently reside in 28 countries whose differences in policy could complicate range-wide management (61) and act as additional artificial barriers. Cooperative international management would be needed to restore historical levels of connectedness. Currently, the African Lion Working Group recommends using regional guidelines for sourcing lions for translocations (62). Though mtDNA structure should still be considered, strict guidelines dictated by nDNA genetic similarities within regional populations may not be as critical for maintaining the population’s genetic diversity if the goal is to reflect historical levels of gene flow.

Connectivity is critical to enable gene flow between subpopulations to avoid the erosion of genetic diversity (63). As an iconic flagship species, these results expose the influence of habitat fragmentation, potentially affecting hundreds of other species. We already are observing the initial effects of this fragmentation on lions through increased nDNA structure and decreased nDNA diversity. Science-based management policies and informed stewardship can help mitigate the loss of nDNA diversity and continued preservation of mtDNA diversity. Intervention is needed to increase gene flow between lion subpopulations and reduce the effects of habitat fragmentation.

## METHODS

### Nuclear Analysis

Biological material from 162 lions dating prior to 1947 was collected from museums (Figure 1, SI Appendix S9) in the form of bone fragments, whole teeth or tooth fragments, nasal turbinate bones, and/or dried tissue. Microsatellite amplification was performed following protocols and procedures described in Curry and Derr 2019 (64). Nine microsatellite loci (Leo006, Leo008, Leo085, Leo098, Leo126, Leo224, Leo230, Leo247, Leo281) that had greater than 75% allele call coverage across both the modern dataset (MD) and the historical nuclear dataset (HD) for all loci were employed in the final analyses. Only lions with known sampling date and location and >70% amplification success were used in downstream analyses. Sample preparation, DNA extraction, and storage, PCR amplification, allele calling, and call verification followed protocols described in Curry and Derr 2019 (64). Further details can be found in SI Appendix S11.

The MD consists of microsatellite allele calls from Bertola *et al*. 2015 (39) (MD-1), Driscoll *et al*. 2002 (65) (MD-2) and Curry *et al*. 2019 (32) (MD-3). Six additional lions were included from the African Wildlife Genomics collection at Texas A&M University (MD-4). These datasets were combined to expand sample size and range for structure analysis and population statistics as well as for direct comparison with the HD. Data calibration is needed when combining microsatellite allele calls from different studies (66). Details on calibration of allele calls can be found in SI Appendix S10.

Nuclear diversity calculations were done using Arlequin v3.5 (67), GenePop (68), HPRare (69), and GenAlEx v6.5 (70). MD and HD were analyzed separately, and results were then compared. A comparison of means was used to determine statistical significance of differences between historical to modern metrics.

Knowing that population structure has been found regionally (28, 30, 33, 37, 40, 71, 72), we implemented a hierarchical strategy to uncover any hidden structure that may be lost when subpopulations are analyzed together (73–75). STRUCTURE runs were performed on each of the full MD and HD datasets without priors for 15 iterations of K 1-15 for 100,000 MCMC reps with 10% burn in. STRUCTURE was rerun for each population as determined through ΔK values from STRUCTURE HARVESTER (76) with individuals assigned to populations based on *Q* scores from runs combined in CLUMPP (77). To determine structural tiers, this was continued until no additional population structure was found. Samples were assigned to the finest level of structure then run as a full population with location priors for 15 iterations of K 1-12. Runs were combined using CLUMPP (77) and visualized at each tier using DISTRUCT (78). To further look at structure patterns, a mantel test for IBD and principal coordinate analysis (PCoA) were performed in GenAlEx v6.5(70).

### Mitochondrial analysis

Only polymorphic sites found in the sequences generated in this study were used for downstream analyses. Conservative filtering was implemented to accommodate the higher error rate associated with DNA damage possible in older samples (79–81). However, conservative filtering in the historical mitogenomes may increase false negative variation present in the published, modern mitogenomes. Therefore, polymorphic sites found only in the published mitogenomes were excluded to reduce potential biases produced by differences in sequencing between studies

Whole mitogenomes were assembled based on whole genome sequencing of 155 samples (152 historical and 3 modern). Details on whole genome sequencing, quality filtering, and SNP identification can be found in SI Appendix S11. After filtering, 102 historical lions and 3 modern lions were of sufficient quality for downstream analyses. Sixteen additional modern lion sequences from GenBank (KP001493-KP001506 (37), KP202262 (82), KC834784 (83)) were added for a total of 19 modern lions.

Mitochondrial diversity analyses of the multiple sequence alignment (MSA) of 280 polymorphic sites were performed using PLINK v1.9 (84), Arlequin v3.5 (67), and DnaSP v6 (85). Principal components analysis (PCA) was performed using R package SNPRelate (86) through calculation of Eigenvectors and visualized using the plot3D function in the rgl R package. A median-joining haplotype network was created using POPArt (87) and an unrooted maximum likelihood (ML) tree was inferred in RAxML using a rapid bootstrap with 1000 replicates evaluated under the GTR+GAMMA+I substitution model (88).

## ACKNOWLEDGEMENTS

We extend our deepest appreciation to the natural history museums that allowed us to sample their collections: American Museum of Natural History, Carnegie Museum of Natural History, Field Museum of Natural History, Kansas University Natural History Museum, Natural History Museum of Los Angeles County, Naturalis Biodiversity Center, Royal Belgian Institute of Natural Sciences, Swedish Royal Museum of Natural History, The Museum of Vertebrate Zoology at Berkeley, Yale Peabody Museum, and Zoological Museum Amsterdam. Additionally, we are grateful to Drs. Hans de Iongh and Klaas Vrieling for their assistance in the procurement of data and samples. A special thank you goes to the laboratory personnel at the DNA Technologies Core Laboratory at Texas A&M University for their assistance during laboratory procedures and analysis. This project was made possible in part by the Texas A&M University College of Veterinary Medicine Trainee Grant, Dallas Safari Club, Dallas Safari Club Foundation, Safari Club International Foundation, Houston Safari Club Foundation Dan L. Duncan Scholarship Award Program, the Explorer’s Club Exploration Fund, the Boore Family Foundation, Curry Family donations, and all the generous backers of the Experiment.com Cat Challenge.

