## Supplementary material for "Spatiotemporal Genetic Diversity of Lions": SI Appendix

**TABLE OF CONTENTS**

S1. Sample Lists
a: Modern Lions – Nuclear Analysis (2 Pages)
b: Modern Lions – Mitochondrial Analysis
c: Historical Lions – Location Information (2 Pages)
d: Historical Lions – Results (2 Pages)

S2. Step-by-step Hierarchical STRUCTURE

S3. Modern Dataset Diversity Statistics

S4. Additional HD PCoA Analyses

S5. Mantel Tests for Isolation-by-Distance

S6. Mitogenome PCA

S7. Mitogenome Haplotype Network

S8. Comparison of Mitochondrial Clustering Across Studies

S9. Table of Museums

S10. Additional Methods

Table 1: STR Calibration Results
Figure 1: Mitogenome Alignments with and without NUMT Correction

S11. Annotation of Lion Mitogenome

S1.a: Modern Lions – Nuclear Analysis (2 Pages)

| | Alternate IDs | | | Sampling | Hierarchical STRUCTURE Results | | | | | |
|---|---|---|---|---|---|---|---|---|---|---|
| Project ID | MD-1 | MD-2 | MD-3 | Region | Country | Location | Continental | Subcontinental | Regional | Local |
| M_GIR_001 | Ple50 | India7 | | Northern | GIR | Gir Forest NP | Asia | Northern | India | GIR |
| M_GIR_002 | Ple53 | India5 | | Northern | GIR | Gir Forest NP | Asia | Northern | India | GIR |
| M_GIR_003 | Ple58 | India2 | | Northern | GIR | Gir Forest NP | Asia | Northern | India | GIR |
| M_GIR_004 | Ple60 | India10 | | Northern | GIR | Gir Forest NP | Asia | Northern | India | GIR |
| M_GIR_005 | Ple67 | India3 | | Northern | GIR | Gir Forest NP | Asia | Northern | India | GIR |
| M_GIR_006 | Ple68 | India1 | | Northern | GIR | Gir Forest NP | Asia | Northern | India | GIR |
| M_GIR_007 | Ple69 | India8 | | Northern | GIR | Gir Forest NP | Asia | Northern | India | GIR |
| M_GIR_008 | Ple70 | India9 | | Northern | GIR | Gir Forest NP | Asia | Northern | India | GIR |
| M_GIR_009 | Ple71 | India6 | | Northern | GIR | Gir Forest NP | Asia | Northern | India | GIR |
| M_GIR_010 | Ple72 | India4 | | Northern | GIR | Gir Forest NP | Asia | Northern | India | GIR |
| M_BEN_011 | | Benin01 | | Western | BEN | Pendjari NP | Africa | Western | West | WES |
| M_BEN_012 | | Benin02 | | Western | BEN | Pendjari NP | Africa | Western | West | WES |
| M_BEN_013 | | Benin03 | | Western | BEN | Pendjari NP | Africa | Western | West | WES |
| M_BEN_014 | | Benin04 | | Western | BEN | Pendjari NP | Africa | Western | West | WES |
| M_BEN_015 | | Benin05 | | Western | BEN | Pendjari NP | Africa | Western | West | WES |
| M_CAM_025 | | Cameroon10 | | Western | CAM | Bénoué Ecosystem | Africa | Western | West | CEN |
| M_CAM_026 | | Cameroon11 | | Western | CAM | Bénoué Ecosystem | Africa | Western | West | CEN |
| M_CAM_027 | | Cameroon12 | | Western | CAM | Bénoué Ecosystem | Africa | Western | West | CEN |
| M_CAM_016 | | Cameroon01 | | Western | CAM | Waza NP | Africa | Western | West | MID |
| M_CAM_017 | | Cameroon02 | | Western | CAM | Waza NP | Africa | Western | West | MID |
| M_CAM_018 | | Cameroon03 | | Western | CAM | Waza NP | Africa | Western | West | MID |
| M_CAM_019 | | Cameroon04 | | Western | CAM | Waza NP | Africa | Western | West | MID |
| M_CAM_020 | | Cameroon05 | | Western | CAM | Waza NP | Africa | Western | West | MID |
| M_CAM_021 | | Cameroon06 | | Western | CAM | Waza NP | Africa | Western | West | MID |
| M_CAM_022 | | Cameroon07 | | Western | CAM | Waza NP | Africa | Western | West | MID |
| M_CAM_023 | | Cameroon08 | | Western | CAM | Waza NP | Africa | Western | West | MID |
| M_CAM_024 | | Cameroon09 | | Western | CAM | Waza NP | Africa | Western | West | MID |
| M_CHD_028 | | Chad01 | | Western | CHD | Zakouma NP | Africa | Western | West | CEN |
| M_CHD_029 | | Chad02 | | Western | CHD | Zakouma NP | Africa | Western | West | CEN |
| M_CHD_030 | | Chad03 | | Western | CHD | Zakouma NP | Africa | Western | West | CEN |
| M_CHD_031 | | Chad04 | | Western | CHD | Zakouma NP | Africa | Western | West | CEN |
| M_DRC_032 | | DRC01 | | Western | DRC | Garamba NP | Africa | Western | West | CEN |
| M_DRC_033 | | DRC02 | | Western | DRC | Garamba NP | Africa | Western | West | CEN |
| M_DRC_034 | | DRC04 | | Western | DRC | Garamba NP | Africa | Western | West | CEN |
| M_DRC_035 | | DRC06 | | Western | DRC | Garamba NP | Africa | Western | West | CEN |
| M_DRC_132 | | DRC03 | | Western | DRC | Garamba NP | Africa | Southern | Southeast | UA |
| M_DRC_133 | | DRC05 | | Western | DRC | Garamba NP | Africa | UA | UA | UA |
| M_DRC_134 | | DRC07 | | Western | DRC | Garamba NP | Africa | UA | UA | UA |
| M_KEN_037 | | Kenya01 | | Eastern | KEN | Amboseli NP | Africa | Eastern | East | KEN |
| M_KEN_038 | | Kenya02 | | Eastern | KEN | Amboseli NP | Africa | Eastern | East | KEN |
| M_KEN_039 | | Kenya03 | | Eastern | KEN | Amboseli NP | Africa | Eastern | East | KEN |
| M_KEN_040 | | Kenya04 | | Eastern | KEN | Amboseli NP | Africa | Eastern | East | KEN |
| M_KEN_041 | | Kenya05 | | Eastern | KEN | Amboseli NP | Africa | Eastern | East | KEN |
| M_KEN_042 | | Kenya06 | | Eastern | KEN | Amboseli NP | Africa | Eastern | East | KEN |
| M_KEN_043 | | Kenya07 | | Eastern | KEN | Amboseli NP | Africa | Eastern | East | KEN |
| M_TAN_044 | Ple235 | Tanzania8 | | Eastern | TAN | Ngorongoro CA | Africa | Eastern | East | TAN |
| M_TAN_045 | Ple239 | Tanzania3 | | Eastern | TAN | Ngorongoro CA | Africa | Eastern | East | TAN |
| M_TAN_056 | Ple349 | Tanzania2 | | Eastern | TAN | Ngorongoro CA | Africa | Eastern | East | TAN |
| M_TAN_057 | Ple359 | Tanzania1 | | Eastern | TAN | Ngorongoro CA | Africa | Eastern | East | TAN |
| M_TAN_058 | Ple490 | Tanzania6 | | Eastern | TAN | Ngorongoro CA | Africa | Eastern | East | TAN |
| M_TAN_059 | Ple493 | Tanzania10 | | Eastern | TAN | Ngorongoro CA | Africa | Eastern | East | TAN |
| M_TAN_060 | Ple508 | Tanzania5 | | Eastern | TAN | Ngorongoro CA | Africa | Eastern | East | TAN |
| M_TAN_061 | Ple510 | Tanzania7 | | Eastern | TAN | Ngorongoro CA | Africa | Eastern | East | TAN |
| M_TAN_062 | Ple539 | Tanzania4 | | Eastern | TAN | Ngorongoro CA | Africa | Eastern | East | TAN |
| M_TAN_063 | Ple572 | Tanzania9 | | Eastern | TAN | Ngorongoro CA | Africa | Eastern | East | TAN |
| M_TAN_046 | Ple276 | Tanzania19 | | Eastern | TAN | Serengeti NP | Africa | Eastern | East | TAN |
| M_TAN_047 | Ple279 | Tanzania12 | | Eastern | TAN | Serengeti NP | Africa | Eastern | East | TAN |
| M_TAN_048 | Ple289 | Tanzania20 | | Eastern | TAN | Serengeti NP | Africa | Eastern | East | TAN |
| M_TAN_049 | Ple301 | Tanzania11 | | Eastern | TAN | Serengeti NP | Africa | Eastern | East | TAN |
| M_TAN_050 | Ple305 | Tanzania13 | | Eastern | TAN | Serengeti NP | Africa | Eastern | East | TAN |
| M_TAN_051 | Ple307 | Tanzania14 | | Eastern | TAN | Serengeti NP | Africa | Eastern | East | TAN |
| M_TAN_052 | Ple311 | Tanzania15 | | Eastern | TAN | Serengeti NP | Africa | Eastern | East | TAN |
| M_TAN_053 | Ple313 | Tanzania16 | | Eastern | TAN | Serengeti NP | Africa | Eastern | East | TAN |
| M_TAN_054 | Ple315 | Tanzania17 | | Eastern | TAN | Serengeti NP | Africa | Eastern | East | TAN |
| M_TAN_055 | Ple317 | Tanzania18 | | Eastern | TAN | Serengeti NP | Africa | Eastern | East | TAN |
| M_TAN_036 | | | | Eastern | TAN | Tanzania | Africa | Western | West | CEN |
| M_TAN_135 | | | | Eastern | TAN | Tanzania | Africa | Southern | Southeast | UA |
| M_ZAM_078 | | Zambia9 | 2011000317 | Southern | ZAM | Kafue NP | Africa | Southern | Southwest | ZAW |

S1.a: Modern Lions – Nuclear Analysis (Page 2 of 2)

| | Alternate IDs | | | Sampling | | | Hierarchical STRUCTURE Results | | | |
|---|---|---|---|---|---|---|---|---|---|---|
| Project ID | MD-1 | MD-2 | MD-3 | Region | Country | Location | Continental | Subcontinental | Regional | Local |
| M_ZAM_079 | | | 2011000325 | Southern | ZAM | Kafue NP | Africa | Southern | Southwest | ZAW |
| M_ZAM_080 | | | 2011000397 | Southern | ZAM | Kafue NP | Africa | Southern | Southwest | ZAW |
| M_ZAM_081 | | | 2011000481 | Southern | ZAM | Kafue NP | Africa | Southern | Southwest | ZAW |
| M_ZAM_082 | | | 2011000482 | Southern | ZAM | Kafue NP | Africa | Southern | Southwest | ZAW |
| M_ZAM_083 | | | 2011000738 | Southern | ZAM | Kafue NP | Africa | Southern | Southwest | ZAW |
| M_ZAM_084 | | | 2011000740 | Southern | ZAM | Kafue NP | Africa | Southern | Southwest | ZAW |
| M_ZAM_085 | | | 2011000746 | Southern | ZAM | Kafue NP | Africa | Southern | Southwest | ZAW |
| M_ZAM_086 | | | 2011000748 | Southern | ZAM | Kafue NP | Africa | Southern | Southwest | ZAW |
| M_ZAM_087 | | | 2011000763 | Southern | ZAM | Kafue NP | Africa | Southern | Southwest | ZAW |
| M_ZAM_088 | | | 2011000784 | Southern | ZAM | Kafue NP | Africa | Southern | Southwest | ZAW |
| M_ZAM_089 | | | 2011000817 | Southern | ZAM | Kafue NP | Africa | Southern | Southwest | ZAW |
| M_ZAM_090 | | | 2011000819 | Southern | ZAM | Kafue NP | Africa | Southern | Southwest | ZAW |
| M_ZAM_115 | | | 2011000310 | Southern | ZAM | Kafue NP | Africa | Southern | Southwest | ETO |
| M_ZAM_128 | | | 2011000401 | Southern | ZAM | Kafue NP | Africa | Southern | UA | UA |
| M_ZAM_129 | | | 2011000409 | Southern | ZAM | Kafue NP | Africa | Southern | UA | UA |
| M_ZAM_131 | | | 2011000883 | Southern | ZAM | Kafue NP | Africa | Southern | UA | UA |
| M_ZAM_064 | | | 2011000286 | Southern | ZAM | Luangwa Valley | Africa | Southern | Southeast | ZAE |
| M_ZAM_065 | | Zambia5 | 2011000298 | Southern | ZAM | Luangwa Valley | Africa | Southern | Southeast | ZAE |
| M_ZAM_066 | | Zambia6 | 2011000299 | Southern | ZAM | Luangwa Valley | Africa | Southern | Southeast | ZAE |
| M_ZAM_067 | | Zambia2 | 2011000300 | Southern | ZAM | Luangwa Valley | Africa | Southern | Southeast | ZAE |
| M_ZAM_068 | | Zambia3 | 2011000301 | Southern | ZAM | Luangwa Valley | Africa | Southern | Southeast | ZAE |
| M_ZAM_069 | | Zambia1 | 2011000302 | Southern | ZAM | Luangwa Valley | Africa | Southern | Southeast | ZAE |
| M_ZAM_070 | | Zambia4 | 2011000305 | Southern | ZAM | Luangwa Valley | Africa | Southern | Southeast | ZAE |
| M_ZAM_071 | | | 2011000312 | Southern | ZAM | Luangwa Valley | Africa | Southern | Southeast | ZAE |
| M_ZAM_072 | | Zambia7 | 2011000324 | Southern | ZAM | Luangwa Valley | Africa | Southern | Southeast | ZAE |
| M_ZAM_073 | | Zambia8 | 2011000353 | Southern | ZAM | Luangwa Valley | Africa | Southern | Southeast | ZAE |
| M_ZAM_074 | | | 2011000418 | Southern | ZAM | Luangwa Valley | Africa | Southern | Southeast | ZAE |
| M_ZAM_075 | | | 2011000699 | Southern | ZAM | Luangwa Valley | Africa | Southern | Southeast | ZAE |
| M_ZAM_076 | | | 2011000706 | Southern | ZAM | Luangwa Valley | Africa | Southern | Southeast | ZAE |
| M_ZAM_077 | | | 2011000849 | Southern | ZAM | Luangwa Valley | Africa | Southern | Southeast | ZAE |
| M_ZAM_094 | | | 2011000881 | Southern | ZAM | Luangwa Valley | Africa | Southern | Southeast | KRU |
| M_ZAM_126 | | | 2011000288 | Southern | ZAM | Luangwa Valley | Africa | Southern | Southeast | UA |
| M_ZAM_127 | | | 2011000399 | Southern | ZAM | Luangwa Valley | Africa | Southern | Southeast | UA |
| M_NAM_116 | Ple402 | Namibia1 | | Southern | NAM | Etoshia NP | Africa | Southern | Southwest | ETO |
| M_NAM_117 | Ple406 | Namibia2 | | Southern | NAM | Etoshia NP | Africa | Southern | Southwest | ETO |
| M_NAM_118 | Ple409 | Namibia3 | | Southern | NAM | Etoshia NP | Africa | Southern | Southwest | ETO |
| M_NAM_119 | Ple411 | Namibia4 | | Southern | NAM | Etoshia NP | Africa | Southern | Southwest | ETO |
| M_NAM_120 | Ple425 | Namibia5 | | Southern | NAM | Etoshia NP | Africa | Southern | Southwest | ETO |
| M_NAM_121 | Ple430 | Namibia6 | | Southern | NAM | Etoshia NP | Africa | Southern | Southwest | ETO |
| M_NAM_122 | Ple437 | Namibia8 | | Southern | NAM | Etoshia NP | Africa | Southern | Southwest | ETO |
| M_NAM_123 | Ple439 | Namibia9 | | Southern | NAM | Etoshia NP | Africa | Southern | Southwest | ETO |
| M_NAM_124 | Ple443 | Namibia7 | | Southern | NAM | Etoshia NP | Africa | Southern | Southwest | ETO |
| M_NAM_125 | Ple448 | Namibia10 | | Southern | NAM | Etoshia NP | Africa | Southern | Southwest | ETO |
| M_RSA_105 | Ple701 | RSA1 | | Southern | RSA | Kalahari-Gemsbok NP | Africa | Southern | Southwest | KAL |
| M_RSA_106 | Ple705 | RSA2 | | Southern | RSA | Kalahari-Gemsbok NP | Africa | Southern | Southwest | KAL |
| M_RSA_107 | Ple707 | RSA3 | | Southern | RSA | Kalahari-Gemsbok NP | Africa | Southern | Southwest | KAL |
| M_RSA_108 | Ple708 | RSA4 | | Southern | RSA | Kalahari-Gemsbok NP | Africa | Southern | Southwest | KAL |
| M_RSA_109 | Ple710 | RSA5 | | Southern | RSA | Kalahari-Gemsbok NP | Africa | Southern | Southwest | KAL |
| M_RSA_110 | Ple711 | RSA6 | | Southern | RSA | Kalahari-Gemsbok NP | Africa | Southern | Southwest | KAL |
| M_RSA_111 | Ple713 | RSA7 | | Southern | RSA | Kalahari-Gemsbok NP | Africa | Southern | Southwest | KAL |
| M_RSA_112 | Ple720 | RSA8 | | Southern | RSA | Kalahari-Gemsbok NP | Africa | Southern | Southwest | KAL |
| M_RSA_113 | Ple721 | RSA9 | | Southern | RSA | Kalahari-Gemsbok NP | Africa | Southern | Southwest | KAL |
| M_RSA_114 | Ple724 | RSA10 | | Southern | RSA | Kalahari-Gemsbok NP | Africa | Southern | Southwest | KAL |
| M_RSA_095 | Ple150 | RSA11 | | Southern | RSA | Kruger NP | Africa | Southern | South | KRU |
| M_RSA_096 | Ple154 | RSA12 | | Southern | RSA | Kruger NP | Africa | Southern | South | KRU |
| M_RSA_097 | Ple155 | RSA13 | | Southern | RSA | Kruger NP | Africa | Southern | South | KRU |
| M_RSA_098 | Ple157 | RSA14 | | Southern | RSA | Kruger NP | Africa | Southern | South | KRU |
| M_RSA_099 | Ple165 | RSA15 | | Southern | RSA | Kruger NP | Africa | Southern | South | KRU |
| M_RSA_100 | Ple168 | RSA16 | | Southern | RSA | Kruger NP | Africa | Southern | South | KRU |
| M_RSA_101 | Ple171 | RSA17 | | Southern | RSA | Kruger NP | Africa | Southern | South | KRU |
| M_RSA_102 | Ple174 | RSA18 | | Southern | RSA | Kruger NP | Africa | Southern | South | KRU |
| M_RSA_103 | Ple179 | RSA19 | | Southern | RSA | Kruger NP | Africa | Southern | South | KRU |
| M_RSA_104 | Ple181b | RSA20 | | Southern | RSA | Kruger NP | Africa | Southern | South | KRU |
| M_RSA_091 | | | | Southern | RSA | South Africa | Africa | Southern | Southeast | KRU |
| M_RSA_092 | | | | Southern | RSA | South Africa | Africa | Southern | Southeast | KRU |
| M_RSA_093 | | | | Southern | RSA | South Africa | Africa | Southern | Southeast | KRU |
| M_RSA_130 | | | | Southern | RSA | South Africa | Africa | Southern | UA | UA |

### S1.b: Modern Lions – Mitochondrial Analysis

| | | | | Sampling | | | Original | | NGS Coverage | | mtDNA Sequence Variation | | | | |
|---|---|---|---|---|---|---|---|---|---|---|---|---|---|---|---|
| Project ID | Accession # | Haplotype | Clade | Region | Country | Location | Material | Protocol | DNA | mtDNA | % Called | %Missing | REF | ALT | ? |
| M_ZAM_T256 | This study | Hap_39 | M | Southern | ZAM | Kafue | Hide | Tissue | 1.7 | 72.0 | 99.9% | 0.1% | 1811 | 0 | 1 |
| M_ZAM_T429 | This study | Hap_39 | M | Southern | ZAM | Kafue | Hide | Tissue | 1.3 | 25.1 | 98.1% | 1.9% | 1777 | 0 | 35 |
| M_TAN_135 | This study | Hap_37 | M | Eastern | TAN | Tanzania | Bone | Bone | 1.6 | 331.0 | 99.9% | 0.1% | 1790 | 21 | 1 |
| M_BEN_011 | KP001497 | Hap_8 | W | Western | BEN | Pendjari NP | | | | | | | | | |
| M_CAM_020 | KP001502 | Hap_21 | W | Western | CAM | Waza NP | | | | | | | | | |
| M_CAM_025 | KP001493 | Hap_22 | W | Western | CAM | Bénoué Ecosystem | | | | | | | | | |
| M_DRC_033 | KP001506 | Hap_17 | W | Western | DRC | Garamba NP | | | | | | | | | |
| M_DRC_KP494 | KP001494 | Hap_20 | W | Western | DRC | Captive (i) | | | | | | | | | |
| M_ETH_KP495 | KP001495 | Hap_23 | W | Eastern | ETH | Ethiopia | | | | | | | | | |
| M_GIR_009 | KP001501 | Hap_6 | W | Northern | GIR | Gir Forest NP | | | | | | | | | |
| M_GIR_KC784 | KC834784 | Hap_5 | W | Northern | GIR | India | | | | | | | | | |
| M_KEN_KP498 | KP001498 | Hap_66 | M | Eastern | KEN | Tsavo East NP | | | | | | | | | |
| M_NAM_KP496 | KP001496 | Hap_74 | S | Southern | NAM | Captive | | | | | | | | | |
| M_NAM_KP504 | KP001504 | Hap_86 | S | Southern | NAM | Eastern Etosha | | | | | | | | | |
| M_PLE_KP262 | KP202262 | Hap_19 | W | Western | UNK | Captive | | | | | | | | | |
| M_RSA_KP500 | KP001500 | Hap_71 | S | Southern | RSA | Kruger NP: Timbavati | | | | | | | | | |
| M_SOM_KP499 | KP001499 | Hap_33 | E | Eastern | SOM | Captive (vi) | | | | | | | | | |
| M_ZAM_KP503 | KP001503 | Hap_38 | M | Southern | ZAM | Mpika town | | | | | | | | | |
| M_ZAM_KP505 | KP001505 | Hap_39 | M | Southern | ZAM | Mulobezi town | | | | | | | | | |

S1.c: Historical Lions – Location Information (2 Pages) – Dates in italics were extrapolated from expedition.

| Project ID | Museum ID | Sampling Region | Current Country | Year | Latitude | Longitude | Location (from Museum Record) | Collector/Expedition |
|---|---|---|---|---|---|---|---|---|
| H_BOT_001 | AMNH_119594 | Southern | BOT | *1930* | -19.4040 | 23.5040 | Ngamiland | Vernay-Lang Kalahari Expedition |
| H_BOT_002 | AMNH_119595 | Southern | BOT | *1930* | -19.4040 | 23.5040 | Ngamiland | Vernay-Lang Kalahari Expedition |
| H_COG_003 | AMNH_119870 | Western | COG | *1949* | -4.2548 | 15.2518 | French Equetorial Africa, Environsde Brazzaville | MacLatchy-Malbrant |
| H_GAB_004 | AMNH_119871 | Western | GAB | *1949* | -0.0925 | 11.9486 | French Equetorial Africa, Booue Gabon | MacLatchy-Malbrant |
| H_CPT_005 | AMNH_13904 | Captive | CAP | 1898 | | | Captive | |
| H_CPT_006 | AMNH_13998 | Captive | CAP | 1898 | | | Captive | |
| H_CPT_007 | AMNH_14027 | Captive | CAP | 1895 | | | Captive | |
| H_CPT_008 | AMNH_14028 | Captive | CAP | 1895 | | | Captive | |
| H_CPT_009 | AMNH_14034 | Captive | CAP | 1896 | | | Captive | |
| H_ZIM_010 | AMNH_161011 | Southern | ZAM | 1946 | -14.0000 | 27.0000 | Rhodesia: Namwaka | T.D. Carter |
| H_MWI_011 | AMNH_161732 | Southern | MWI | 1946 | -13.2734 | 33.6373 | Nyasaland: Ntchisi to Chibotela | H.E. Anthony |
| H_COG_012 | AMNH_17274 | Western | COG | *<1920* | -4.2548 | 15.2518 | Congo Region | Mr. Glove |
| H_COG_013 | AMNH_17275 | Western | COG | *<1920* | -4.2548 | 15.2518 | Congo Region | Mr. Glove, Gift of the Century Co. |
| H_NAM_014 | AMNH_19181 | Southern | NAM | *<1913* | -19.0857 | 16.3993 | German-South-West Africa | H.H. Vogelsang Collection (2798/1211) |
| H_CPT_015 | AMNH_24249 | Captive | CAP | 1905 | | | Captive | Original No. 1440 |
| H_KEN_016 | AMNH_27769 | Eastern | KEN | 1906 | -0.1000 | 36.1000 | British East Africa | Tjader Expedition |
| H_RSA_017 | AMNH_28151 | Southern | RSA | *1905* | -29.0000 | 24.0000 | South Africa | Richard Douglas Collection |
| H_KEN_018 | AMNH_30240 | Eastern | KEN | *1912* | -0.1000 | 36.1000 | British East Africa | Paul J. Rainey Expedition |
| H_KEN_019 | AMNH_30241 | Eastern | KEN | *1912* | -0.1000 | 36.1000 | British East Africa | Paul J. Rainey Expedition |
| H_KEN_020 | AMNH_30242 | Eastern | KEN | *1912* | -0.1000 | 36.1000 | British East Africa | Paul J. Rainey Expedition |
| H_KEN_021 | AMNH_30243 | Eastern | KEN | *1912* | -0.1000 | 36.1000 | British East Africa | Paul J. Rainey Expedition |
| H_KEN_022 | AMNH_30244 | Eastern | KEN | *1912* | -0.1000 | 36.1000 | British East Africa | Paul J. Rainey Expedition |
| H_KEN_023 | AMNH_30245 | Eastern | KEN | *1912* | -0.1000 | 36.1000 | British East Africa | Paul J. Rainey Expedition |
| H_KEN_024 | AMNH_30246 | Eastern | KEN | *1912* | -0.1000 | 36.1000 | British East Africa | Paul J. Rainey Expedition |
| H_KEN_025 | AMNH_30247 | Eastern | KEN | *1912* | -0.1000 | 36.1000 | British East Africa | Paul J. Rainey Expedition |
| H_KEN_026 | AMNH_30248 | Eastern | KEN | *1912* | -0.1000 | 36.1000 | British East Africa | Paul J. Rainey Expedition |
| H_CPT_027 | AMNH_35472 | Captive | CAP | 1912 | | | Captive | |
| H_KEN_028 | AMNH_36420 | Eastern | KEN | *1912* | -0.1000 | 36.1000 | British East Africa | Paul J. Rainey Expedition |
| H_KEN_029 | AMNH_36421 | Eastern | KEN | *1912* | -0.1000 | 36.1000 | British East Africa | Paul J. Rainey Expedition |
| H_CPT_030 | AMNH_403 | Captive | CAP | 1886 | | | Captive | |
| H_DRC_031 | AMNH_52070 | Western | DRC | 1911 | 3.7327 | 29.7164 | Belgian Congo: Faradje | American Museum Congo Expedition |
| H_DRC_032 | AMNH_52071 | Western | DRC | 1911 | 3.7327 | 29.7164 | Belgian Congo: Faradje | American Museum Congo Expedition |
| H_DRC_033 | AMNH_52072 | Western | DRC | 1911 | 3.7327 | 29.7164 | Belgian Congo: Faradje | American Museum Congo Expedition |
| H_DRC_034 | AMNH_52073 | Western | DRC | 1911 | 3.7327 | 29.7164 | Belgian Congo: Faradje | American Museum Congo Expedition |
| H_DRC_035 | AMNH_52074 | Western | DRC | 1911 | 3.7327 | 29.7164 | Belgian Congo: Faradje | American Museum Congo Expedition |
| H_DRC_036 | AMNH_52075 | Western | DRC | 1911 | 3.7327 | 29.7164 | Belgian Congo: Faradje | American Museum Congo Expedition |
| H_DRC_037 | AMNH_52076 | Western | DRC | 1911 | 3.7327 | 29.7164 | Belgian Congo: Faradje | American Museum Congo Expedition |
| H_DRC_038 | AMNH_52077 | Western | DRC | 1912 | 3.7327 | 29.7164 | Belgian Congo: Faradje | American Museum Congo Expedition |
| H_DRC_039 | AMNH_52078 | Western | DRC | 1912 | 3.7327 | 29.7164 | Belgian Congo: Faradje | American Museum Congo Expedition |
| H_DRC_040 | AMNH_52079 | Western | DRC | 1912 | 3.7327 | 29.7164 | Belgian Congo: Faradje | American Museum Congo Expedition |
| H_DRC_041 | AMNH_52080 | Western | DRC | 1912 | 3.7327 | 29.7164 | Belgian Congo: Faradje | American Museum Congo Expedition |
| H_DRC_042 | AMNH_52081 | Western | DRC | 1912 | 3.7327 | 29.7164 | Belgian Congo: Faradje | American Museum Congo Expedition |
| H_DRC_043 | AMNH_52082 | Western | DRC | 1912 | 3.7327 | 29.7164 | Belgian Congo: Faradje | American Museum Congo Expedition |
| H_KEN_044 | AMNH_54370 | Eastern | KEN | 1912 | 0.5160 | 35.3234 | Kenya | Akeley Expedition to British East Africa |
| H_KEN_045 | AMNH_54371 | Eastern | KEN | 1912 | 0.5160 | 35.3234 | Kenya | Akeley Expedition to British East Africa |
| H_KEN_046 | AMNH_54372 | Eastern | KEN | 1912 | 0.5160 | 35.3234 | Kenya | Akeley Expedition to British East Africa |
| H_KEN_047 | AMNH_54393 | Eastern | KEN | 1912 | 0.5160 | 35.3234 | Kenya: Gyasiu Nghishu Plateau | Akeley Expedition to British East Africa |
| H_KEN_048 | AMNH_54394 | Eastern | KEN | 1912 | 0.5160 | 35.3234 | Kenya | Akeley Expedition to British East Africa |
| H_KEN_049 | AMNH_54395 | Eastern | KEN | 1912 | 0.5160 | 35.3234 | Kenya: Gyasiu Nghishu Plateau | Akeley Expedition to British East Africa |
| H_GIR_050 | AMNH_54995 | Northern | GIR | 1929 | 21.1240 | 70.8217 | India: Junagadh State, Gir Forest | Faunthorpe-Vernay Expedition |
| H_GIR_051 | AMNH_54996 | Northern | GIR | 1929 | 21.1240 | 70.8217 | India: Junagadh State, Gir Forest | Faunthorpe-Vernay Expedition |
| H_CPT_052 | AMNH_6260 | Captive | CAP | 1893 | | | Captive | |
| H_CPT_053 | AMNH_6282 | Captive | CAP | 1889 | | | Captive | |
| H_GIR_054 | AMNH_63955 | Northern | GIR | 1906 | 21.1240 | 70.8217 | India | |
| H_CPT_055 | AMNH_65 | Captive | CAP | *1860* | | | Captive | Maximillian Collection |
| H_CPT_056 | AMNH_70171 | Captive | CAP | 1924 | | | Captive | |
| H_KEN_057 | AMNH_70347 | Eastern | KEN | 1922 | -0.1000 | 36.1000 | British East Africa | Wild caught: Donated to NY Zoo by Buffalo Jones |
| H_ANG_058 | AMNH_80609 | Southern | ANG | 1925 | -14.4699 | 16.2911 | Angola: Capelongo | Vernay Angola Expedition/Lang Collection |
| H_RSA_059 | AMNH_81836 | Southern | RSA | *1930* | -25.0000 | 29.0000 | Transvaal | Lang-deLaporte/Lang Collection |
| H_RSA_060 | AMNH_81837 | Southern | RSA | *1930* | -25.0000 | 29.0000 | Transvaal | Lang-deLaporte/Lang Collection |
| H_RSA_061 | AMNH_81839 | Southern | RSA | *1930* | -25.0000 | 29.0000 | Transvaal | Lang-deLaporte/Lang Collection |
| H_RSA_062 | AMNH_81840 | Southern | RSA | *1930* | -25.0000 | 29.0000 | Transvaal | Lang-deLaporte/Lang Collection |
| H_RSA_063 | AMNH_81841 | Southern | RSA | *1930* | -25.0000 | 29.0000 | Transvaal | Lang-deLaporte/Lang Collection |
| H_RSA_064 | AMNH_81842 | Southern | RSA | *1930* | -25.0000 | 29.0000 | Transvaal | Lang-deLaporte/Lang Collection |
| H_RSA_065 | AMNH_81843 | Southern | RSA | *1930* | -25.0000 | 29.0000 | Transvaal | Lang-deLaporte/Lang Collection |
| H_RSA_066 | AMNH_81844 | Southern | RSA | *1930* | -25.0000 | 29.0000 | Transvaal | Lang-deLaporte/Lang Collection |
| H_CAR_067 | AMNH_83410 | Western | CAR | 1924 | 6.9846 | 18.9116 | French Equatorial Africa, NE Section, Ubongui-Churi | Mrs. Martha Bliven |
| H_CPT_068 | AMNH_8355 | Captive | CAP | 1895 | | | Captive | |
| H_BOT_069 | AMNH_83617 | Southern | BOT | *1930* | -19.1492 | 23.7915 | Kalahari | Vernay-Lang Kalahari Expedition |
| H_BOT_070 | AMNH_83618 | Southern | BOT | *1930* | -19.1492 | 23.7915 | Bechuanaland Protectorate: Kwaai, R. Machaba | Vernay-Lang Kalahari Expedition |
| H_BOT_071 | AMNH_83619 | Southern | BOT | *1930* | -19.1492 | 23.7915 | Bechuanaland Protectorate: Kwaai, R. Machaba | Vernay-Lang Kalahari Expedition |
| H_BOT_072 | AMNH_83620 | Southern | BOT | *1930* | -19.1492 | 23.7915 | Bechuanaland Protectorate: Kwaai, R. Machaba | Vernay-Lang Kalahari Expedition |
| H_BOT_073 | AMNH_83621 | Southern | BOT | *1930* | -19.1492 | 23.7915 | Bechuanaland Protectorate: Kwaai, R. Machaba | Vernay-Lang Kalahari Expedition |
| H_BOT_074 | AMNH_83622 | Southern | BOT | *1930* | -19.1492 | 23.7915 | Bechuanaland Protectorate: Kwaai, R. Machaba | Vernay-Lang Kalahari Expedition |
| H_BOT_075 | AMNH_83623 | Southern | BOT | *1930* | -19.1492 | 23.7915 | Bechuanaland Protectorate: Kwaai, R. Machaba | Vernay-Lang Kalahari Expedition |
| H_BOT_076 | AMNH_83624 | Southern | BOT | *1930* | -19.7000 | 26.4000 | Bechuanaland Protectorate: Nkate | Vernay-Lang Kalahari Expedition |
| H_BOT_077 | AMNH_83625 | Southern | BOT | *1930* | -18.7167 | 24.3500 | Bechuanaland Protectorate: Tsotsoroga Pan | Vernay-Lang Kalahari Expedition |
| H_CPT_078 | AMNH_8364 | Captive | CAP | 1895 | | | Captive | |
| H_TAN_079 | AMNH_85140 | Eastern | TAN | 1928 | -6.2582 | 37.7166 | Tanganyika Territory: Serengeti Plains | Carlisle-Clark African Expedition |
| H_TAN_080 | AMNH_85141-L | Eastern | TAN | 1928 | -6.2582 | 37.7166 | Tanganyika Territory: Serengeti Plains | Carlisle-Clark African Expedition |
| H_TAN_081 | AMNH_85141-N | Eastern | TAN | 1928 | -6.2582 | 37.7166 | Tanganyika Territory: Serengeti Plains | Carlisle-Clark African Expedition |
| H_TAN_082 | AMNH_85142-L | Eastern | TAN | 1928 | -6.2582 | 37.7166 | Tanganyika Territory: Serengeti Plains | Carlisle-Clark African Expedition |
| H_TAN_083 | AMNH_85142-N | Eastern | TAN | 1928 | -6.2582 | 37.7166 | Tanganyika Territory: Serengeti Plains | Carlisle-Clark African Expedition |
| H_TAN_084 | AMNH_85143 | Eastern | TAN | 1928 | -6.2582 | 37.7166 | Tanganyika Territory: Serengeti Plains | Carlisle-Clark African Expedition |
| H_TAN_085 | AMNH_85144 | Eastern | TAN | 1928 | -6.2582 | 37.7166 | Tanganyika Territory: Serengeti Plains | Carlisle-Clark African Expedition |
| H_TAN_086 | AMNH_85145 | Eastern | TAN | 1928 | -6.2582 | 37.7166 | Tanganyika Territory: Serengeti Plains | Carlisle-Clark African Expedition |
| H_TAN_087 | AMNH_85146 | Eastern | TAN | 1928 | -6.2582 | 37.7166 | Tanganyika Territory: Serengeti Plains | Carlisle-Clark African Expedition |

S1.c: Historical Lions – Location Information (Page 2 of 2) – Dates in *italics* were extrapolated from expedition.

| Project ID | Museum ID | Sampling Region | Current Country | Year | Latitude | Longitude | Location (from Museum Record) | Collector/Expedition |
|---|---|---|---|---|---|---|---|---|
| H_TAN_088 | AMNH_85147 | Eastern | TAN | 1928 | -6.2582 | 37.7166 | Tanganyika Territory: Serengeti Plains | Carlisle-Clark African Expedition |
| H_TAN_089 | AMNH_85148 | Eastern | TAN | 1928 | -6.2582 | 37.7166 | Tanganyika Territory: Serengeti Plains | Carlisle-Clark African Expedition |
| H_TAN_090 | AMNH_85149 | Eastern | TAN | 1928 | -6.2582 | 37.7166 | Tanganyika Territory: Serengeti Plains | Carlisle-Clark African Expedition |
| H_KEN_091 | AMNH_88632 | Eastern | KEN | *1912* | -0.1000 | 36.1000 | British East Africa | 3rd African Expedition of the American Museum |
| H_KEN_092 | AMNH_88633 | Eastern | KEN | *1912* | -0.1000 | 36.1000 | British East Africa | 3rd African Expedition of the American Museum |
| H_KEN_093 | AMNH_88634 | Eastern | KEN | *1912* | -0.1000 | 36.1000 | British East Africa | 3rd African Expedition of the American Museum |
| H_KEN_094 | AMNH_88635 | Eastern | KEN | *1912* | -0.1000 | 36.1000 | British East Africa | 3rd African Expedition of the American Museum |
| H_KEN_095 | AMNH_88636 | Eastern | KEN | *1912* | -0.1000 | 36.1000 | British East Africa | 3rd African Expedition of the American Museum |
| H_KEN_096 | AMNH_88637 | Eastern | KEN | *1912* | -0.1000 | 36.1000 | British East Africa | 3rd African Expedition of the American Museum |
| H_CPT_097 | CM_1461 | Captive | CAP | | | | Captive | Pittsburgh Zoological Gargen |
| H_CPT_098 | CM_1564 | Captive | CAP | 1908 | | | Captive | Pittsburgh Zoological Gargen |
| H_CPT_099 | CM_1565 | Captive | CAP | 1908 | | | Captive | Pittsburgh Zoological Gargen |
| H_CPT_100 | CM_1825 | Captive | CAP | | | | Captive | Pittsburgh Zoological Gargen |
| H_UNK_101 | CM_184 | Unknown | UNK | | | | | The Klondike Museum |
| H_UNK_102 | CM_185 | Unknown | UNK | | | | | The Klondike Museum |
| H_UNK_103 | CM_31 | Unknown | UNK | | | | | Webster, F S |
| H_ETH_104 | CM_5868 | Eastern | ETH | 1912 | 7.5000 | 40.0000 | Abyssinia, 9-20 MI UP THIKA RIVER* | Frick, C |
| H_TAN_105 | CM_5897 | Eastern | TAN | 1927 | -6.0000 | 35.0000 | Tanzania | Arbuthnot, T H |
| H_TAN_106 | CM_5898 | Eastern | TAN | 1927 | -6.0000 | 35.0000 | Tanzania | Arbuthnot, T H |
| H_TAN_107 | CM_5899 | Eastern | TAN | 1927 | -6.0000 | 35.0000 | Tanzania | Arbuthnot, T H |
| H_TAN_108 | FMNH_127836 | Eastern | TAN | 1928 | -2.4364 | 34.8210 | Mara Region, Bariadi District, Serengeti Plains, Seronera | Cudahy-Massee Expedition |
| H_TAN_109 | FMNH_127837 | Eastern | TAN | 1928 | -2.4364 | 34.8210 | Mara Region, Bariadi District, Serengeti Plains, Seronera | Cudahy-Massee Expedition |
| H_TAN_110 | FMNH_127838 | Eastern | TAN | 1928 | -2.4364 | 34.8210 | Mara Region, Bariadi District, Serengeti Plains, Seronera | Cudahy-Massee Expedition |
| H_TAN_111 | FMNH_127839 | Eastern | TAN | 1928 | -2.4364 | 34.8210 | Mara Region, Bariadi District, Serengeti Plains, Seronera | Cudahy-Massee Expedition |
| H_TAN_112 | FMNH_127840 | Eastern | TAN | | -6.0000 | 35.0000 | Tanzania | |
| H_SOM_113 | FMNH_1443 | Eastern | SOM | 1896 | 10.0000 | 44.0000 | Woqooyi Galbeed, Bannaanko Tuuyo ("Tuyo Plain") | D. G. Elliot |
| H_KEN_114 | FMNH_20756 | Eastern | KEN | 1905 | -1.4613 | 36.9827 | Eastern Prov, Machakos Dist, Athi Plains | Akeley Expedition to British East Africa |
| H_KEN_115 | FMNH_20757 | Eastern | KEN | 1905 | -1.4613 | 36.9827 | Eastern Prov, Machakos Dist, Athi Plains | Akeley Expedition to British East Africa |
| H_KEN_116 | FMNH_20758 | Eastern | KEN | 1905 | -1.4613 | 36.9827 | Eastern Prov, Machakos Dist, Athi Plains | Akeley Expedition to British East Africa |
| H_KEN_117 | FMNH_20760 | Eastern | KEN | 1905 | -1.4613 | 36.9827 | Eastern Prov, Machakos Dist, Athi Plains | Akeley Expedition to British East Africa |
| H_KEN_118 | FMNH_20762 | Eastern | KEN | 1905 | -1.4613 | 36.9827 | Eastern Prov, Machakos Dist, Athi Plains | Akeley Expedition to British East Africa |
| H_SDN_119 | FMNH_30778 | Western | SDN | 1928 | 13.0000 | 35.5000 | Kassala, Rahad R, Abid, Ethiopian border, 20 mi NW | J. E. Baum |
| H_GIR_120 | FMNH_31121 | Northern | GIR | 1929 | 21.1240 | 70.8217 | Kathiawar, Gir Forest | Faunthorpe-Vernay Expedition |
| H_TAN_121 | FMNH_33479 | Eastern | TAN | 1929 | -3.0000 | 35.0000 | Serengeti Plains | C. J. Albrecht |
| H_TAN_122 | FMNH_33480 | Eastern | TAN | 1929 | -3.0000 | 35.0000 | Serengeti Plains | C. J. Albrecht |
| H_TAN_123 | FMNH_35131 | Eastern | TAN | 1930 | -3.0000 | 35.0000 | Serengeti Plains | M. Field |
| H_TAN_124 | FMNH_35132 | Eastern | TAN | 1930 | -3.0000 | 35.0000 | Serengeti Plains | M. Field |
| H_TAN_125 | FMNH_35133 | Eastern | TAN | *1930* | -3.0000 | 35.0000 | Serengeti Plains | M. Field |
| H_TAN_126 | FMNH_35134 | Eastern | TAN | *1930* | -3.0000 | 35.0000 | Serengeti Plains | M. Field |
| H_BOT_127 | FMNH_35739 | Southern | BOT | 1930 | -22.5500 | 23.2500 | Kaotwe Pan, ca 45 mi SE of Gomodimo Pan | Vernay-Lang Kalahari Expedition |
| H_BOT_128 | FMNH_35740 | Southern | BOT | 1930 | -19.1953 | 24.0070 | Mababe Flats | Vernay-Lang Kalahari Expedition |
| H_BOT_129 | FMNH_35741 | Southern | BOT | 1930 | -19.1953 | 24.0070 | Mababe Flats | Vernay-Lang Kalahari Expedition |
| H_BOT_130 | FMNH_35742 | Southern | BOT | 1930 | -19.1953 | 24.0070 | Mababe Flats | Vernay-Lang Kalahari Expedition |
| H_BOT_131 | FMNH_35743 | Southern | BOT | 1930 | -19.7000 | 26.4000 | Nata, 35 mi SW; N'kate | Vernay-Lang Kalahari Expedition |
| H_RSA_132 | FMNH_38134 | Southern | RSA | 1928 | -24.3697 | 30.6441 | Transvaal, Pretoria Dist, Olifants R | H. Lang |
| H_BOT_133 | FMNH_41405 | Southern | BOT | 1930 | -19.1953 | 24.0070 | Mababe Flats | Vernay-Lang Kalahari Expedition |
| H_MLI_134 | FMNH_42129 | Western | MLI | 1920 | 14.3496 | -3.6102 | Mopti, Bandiagara | R. Boulton |
| H_KEN_135 | FMNH_75608 | Eastern | KEN | 1935 | 0.5160 | 35.3234 | Kenya | H. C. Pearson |
| H_KEN_136 | FMNH_75609 | Eastern | KEN | 1935 | 0.5160 | 35.3234 | Kenya | H. C. Pearson |
| H_BOT_137 | FMNH_89926 | Southern | BOT | 1935 | -23.0000 | 24.0000 | Botswana | Vernay-Lang Kalahari Expedition |
| H_SDN_138 | KBIN_469459 | Western | SDN | 1905 | 15.0000 | 30.0000 | Sudan | |
| H_ZAM_139 | KBIN_504512 | Southern | ZAM | 1905 | -8.6606 | 29.8588 | Mweru Wantipa - Zambia | |
| H_TAN_140 | KU_105216 | Eastern | TAN | 1920 | -2.9400 | 37.3400 | Mt Kilomonjaro | Leasure F G |
| H_TAN_141 | KU_105217 | Eastern | TAN | 1920 | -2.9400 | 37.3400 | Mt Kilomonjaro | Leasure F G |
| H_KEN_142 | LACM_51295 | Eastern | KEN | 1922 | -1.3547 | 36.8216 | Nairobi | J. Dines |
| H_KEN_143 | LACM_51296 | Eastern | KEN | 1922 | -1.3547 | 36.8216 | Nairobi | J. Dines |
| H_KEN_144 | LACM_51297 | Eastern | KEN | 1922 | -1.3547 | 36.8216 | Nairobi | J. Dines |
| H_TAN_145 | MVZ_96804 | Eastern | KEN | 1929 | -1.3547 | 36.8216 | Tanganyika District Safari Service from town of Nairobi | Ward C. Russell, A. F. Lieuranse |
| H_ANG_146 | RMNH_45281 | Southern | ANG | 1887 | -15.1996 | 12.1578 | Mossamedes, Angola | |
| H_NRT_147 | RMNH_45282 | Northern | NRT | 1823 | 35.0410 | 0.2941 | Berberleeuw (Barbary: Dutch) | |
| H_MOZ_148 | RMNH_45284 | Southern | MOZ | 1929 | -18.0000 | 35.0000 | Caia omgeving, Mozambique | |
| H_MOZ_149 | RMNH_45285 | Southern | MOZ | 1929 | -18.0000 | 35.0000 | Caia omgeving, Mozambique | |
| H_NAM_150 | S_581971 | Southern | NAM | 1856 | -23.0679 | 14.5063 | Namibia, Erongo, Walvis Bay | |
| H_NRT_151 | S_585287 | Northern | NRT | 1831 | 35.0410 | 0.2941 | Barbariet (Barbary: Swedish) | |
| H_DRC_152 | S_595059 | Western | DRC | 1921 | -0.7556 | 29.3318 | Congo, Ruindi Plains S. of Lake Edward | |
| H_TAN_153 | YPM_2057 | Eastern | TAN | 1928 | -3.0000 | 35.0000 | Tanganyika Territory: Serengeti Plains | S. Clark |
| H_KEN_154 | YPM_3189 | Eastern | KEN | 1931 | -1.0000 | 35.0000 | Rift Valley Province, Masai Reservation, Amorro River | J.C. Rathborne |
| H_KEN_155 | YPM_3217 | Eastern | KEN | 1931 | -1.0000 | 35.0000 | Rift Valley Province, Masai Reservation, Amorro River | J.C. Rathborne |
| H_KEN_156 | YPM_3218 | Eastern | KEN | 1931 | -1.0000 | 35.0000 | Rift Valley Province, Masai Reservation, Amorro River | J.C. Rathborne |
| H_KEN_157 | YPM_5251 | Eastern | KEN | *1922* | -1.0000 | 34.0000 | East Africa | Morden African Expedition |
| H_CPT_158 | YPM_6943 | Captive | CAP | | | | (menagerie) | G.B. Grinnell |
| H_UNK_159 | YPM_6952 | Unknown | UNK | *1917* | | | | H.W. Boyd |
| H_KEN_160 | YPM_9570 | Eastern | KEN | 1931 | -1.0000 | 35.0000 | Rift Valley Province, Masai Reservation, Amorro River | J.C. Rathborne |
| H_RSA_161 | ZMA_11753 | Southern | RSA | 1890 | -24.3697 | 30.6441 | between Pretoria and Lourenco Marques, South Africa | |
| H_CAR_162 | ZMA_996 | Western | CAR | 1905 | 10.2910 | 22.7837 | Birao, Central African Republic | |

### S1.d: Historical Lions – Results (2 Pages)

| | | Datasets | | Original | Extraction | NGS Coverage | | | mtDNA Sequence Variation | | | | | | | STR % Amplification | | |
|---|---|---|---|---|---|---|---|---|---|---|---|---|---|---|---|---|---|---|
| Project ID | Museum ID | nDNA | mtDNA | Material | Protocol | ng/uL | DNA | mtDNA | % Called | %Missing | REF | ALT | ? | Haplotype | Clade | 14 Loci | 8 Loci | 9 Loci |
| H_BOT_001 | AMNH_119594 | X | X | Turbinates | Bone | 17.6 | 2.8 | 27.2 | 96.2% | 3.8% | 1692 | 51 | 69 | Hap_72 | S | 64% | 82% | 82% |
| H_BOT_002 | AMNH_119595 | X | X | Tooth | Bone | 12.6 | 2.2 | 92.7 | 99.6% | 0.4% | 1724 | 80 | 8 | Hap_83 | S | 100% | 100% | 100% |
| H_COG_003 | AMNH_119870 | X | X | Tooth | Bone | 3.1 | 2.1 | 128.6 | 99.7% | 0.3% | 1724 | 82 | 6 | Hap_77 | S | 93% | 100% | 100% |
| H_GAB_004 | AMNH_119871 | X | X | Bone | Bone | 13.2 | 1.8 | 103.0 | 99.7% | 0.3% | 1724 | 82 | 6 | Hap_77 | S | 100% | 100% | 100% |
| H_CPT_005 | AMNH_13904 | | | Bone | Bone | 13.7 | 3.4 | 8.8 | 75.4% | 24.6% | 1357 | 10 | 445 | | | 57% | 73% | 73% |
| H_CPT_006 | AMNH_13998 | | | Bone | Bone | 18.9 | 2.4 | 262.8 | 99.6% | 0.4% | 1785 | 20 | 7 | | | 100% | 100% | 100% |
| H_CPT_007 | AMNH_14027 | | | Turbinates | Bone | 11.8 | 2.9 | 300.8 | 99.6% | 0.4% | 1784 | 20 | 8 | | | 79% | 91% | 91% |
| H_CPT_008 | AMNH_14028 | | | Turbinates | Bone | 8.7 | 3.0 | 665.2 | 99.6% | 0.4% | 1708 | 96 | 8 | | | 43% | 45% | 45% |
| H_CPT_009 | AMNH_14034 | | | Turbinates | Bone | 18.0 | 3.0 | 2.6 | 5.9% | 94.1% | 107 | 0 | 1705 | | | 100% | 100% | 100% |
| H_ZIM_010 | AMNH_161011 | X | X | Bone | Bone | 9.5 | 2.2 | 128.7 | 99.3% | 0.7% | 1724 | 76 | 12 | Hap_76 | S | 100% | 100% | 100% |
| H_MWI_011 | AMNH_161732 | X | X | Tooth | Bone | 1.4 | 2.4 | 85.3 | 99.3% | 0.7% | 1777 | 22 | 13 | Hap_42 | M | 86% | 100% | 100% |
| H_COG_012 | AMNH_17274 | | X | Tooth | Bone | 5.3 | 2.3 | 21.8 | 98.5% | 1.5% | 1763 | 22 | 27 | Hap_40 | M | 64% | 82% | 82% |
| H_COG_013 | AMNH_17275 | | X | Bone | Bone | 14.7 | 2.9 | 103.5 | 99.5% | 0.5% | 1781 | 22 | 9 | Hap_41 | M | 100% | 100% | 100% |
| H_NAM_014 | AMNH_19181 | X | X | Tooth | Bone | 5.9 | 2.0 | 54.4 | 99.0% | 1.0% | 1775 | 19 | 18 | Hap_49 | M | 79% | 91% | 91% |
| H_CPT_015 | AMNH_24249 | | | Tooth | Bone | 3.3 | 2.1 | 172.6 | 99.7% | 0.3% | 1707 | 99 | 6 | | | 86% | 91% | 91% |
| H_KEN_016 | AMNH_27769 | X | X | Hide | Bone | 84.1 | 2.5 | 377.6 | 99.6% | 0.4% | 1725 | 80 | 7 | Hap_32 | E | 79% | 82% | 82% |
| H_RSA_017 | AMNH_28151 | X | | Turbinates | Bone | 5.0 | 3.1 | 7.5 | 39.2% | 60.8% | 700 | 11 | 1101 | | | 71% | 82% | 82% |
| H_KEN_018 | AMNH_30240 | X | | Hide | Bone | 8.9 | 5.4 | 2.0 | 0.1% | 99.9% | 2 | 0 | 1810 | | | 86% | 91% | 91% |
| H_KEN_019 | AMNH_30241 | X | X | Tooth | Bone | 3.3 | 2.3 | 174.0 | 99.5% | 0.5% | 1782 | 21 | 9 | Hap_65 | M | 71% | 82% | 82% |
| H_KEN_020 | AMNH_30242 | X | X | Tooth | Bone | 6.2 | 2.5 | 75.5 | 98.8% | 1.2% | 1769 | 21 | 22 | Hap_65 | M | 93% | 100% | 100% |
| H_KEN_021 | AMNH_30243 | X | X | Tooth | Bone | 11.0 | 2.0 | 47.3 | 98.9% | 1.1% | 1770 | 22 | 20 | Hap_66 | M | 93% | 100% | 100% |
| H_KEN_022 | AMNH_30244 | X | X | Tooth | Bone | 7.1 | 3.1 | 240.2 | 99.6% | 0.4% | 1782 | 22 | 8 | Hap_66 | M | 100% | 100% | 100% |
| H_KEN_023 | AMNH_30245 | X | X | Tooth | Bone | 10.8 | 2.2 | 96.2 | 99.0% | 1.0% | 1772 | 22 | 18 | Hap_66 | M | 93% | 100% | 100% |
| H_KEN_024 | AMNH_30246 | X | X | Tooth | Bone | 1.5 | 3.7 | 202.8 | 98.6% | 1.4% | 1765 | 21 | 26 | Hap_64 | M | 71% | 82% | 82% |
| H_KEN_025 | AMNH_30247 | X | X | Tooth | Bone | 5.6 | 2.4 | 71.2 | 98.8% | 1.2% | 1769 | 22 | 21 | Hap_66 | M | 79% | 100% | 100% |
| H_KEN_026 | AMNH_30248 | X | X | Tooth | Bone | 21.3 | 2.1 | 26.6 | 98.7% | 1.3% | 1765 | 23 | 24 | Hap_43 | M | 100% | 100% | 100% |
| H_CPT_027 | AMNH_35472 | | | Bone | Bone | 51.5 | 3.8 | 3.1 | 11.5% | 88.5% | 205 | 3 | 1604 | | | 93% | 100% | 100% |
| H_KEN_028 | AMNH_36420 | X | X | Tooth | Bone | 13.1 | 2.1 | 169.6 | 99.4% | 0.6% | 1780 | 21 | 11 | Hap_47 | M | 100% | 100% | 100% |
| H_KEN_029 | AMNH_36421 | X | X | Tooth | Bone | 1.0 | 3.5 | 1482.9 | 99.6% | 0.4% | 1783 | 21 | 8 | Hap_47 | M | 71% | 82% | 82% |
| H_CPT_030 | AMNH_403 | | | Bone | Bone | 7.3 | 3.4 | 2.5 | 2.9% | 97.1% | 53 | 0 | 1759 | | | 64% | 64% | 64% |
| H_DRC_031 | AMNH_52070 | X | X | Tooth | Bone | 7.5 | 3.8 | 372.9 | 99.5% | 0.5% | 1709 | 94 | 9 | Hap_13 | W | 93% | 100% | 100% |
| H_DRC_032 | AMNH_52071 | X | | Turbinates | Bone | 149.0 | 4.3 | 10.1 | 81.8% | 18.2% | 1425 | 58 | 329 | | | 93% | 100% | 100% |
| H_DRC_033 | AMNH_52072 | X | X | Bone | Bone | 39.8 | 2.5 | 12.3 | 88.1% | 11.9% | 1536 | 61 | 215 | Hap_7 | W | 93% | 100% | 100% |
| H_DRC_034 | AMNH_52073 | X | X | Bone | Bone | 2.5 | 3.8 | 65.1 | 96.2% | 3.8% | 1723 | 21 | 68 | Hap_51 | M | 93% | 100% | 100% |
| H_DRC_035 | AMNH_52074 | X | X | Tooth | Bone | 2.0 | 2.6 | 35.6 | 98.7% | 1.3% | 1701 | 87 | 24 | Hap_11 | W | 71% | 82% | 82% |
| H_DRC_036 | AMNH_52075 | X | X | Tooth | Bone | 2.9 | 2.6 | 217.8 | 99.6% | 0.4% | 1710 | 95 | 7 | Hap_15 | W | 57% | 64% | 64% |
| H_DRC_037 | AMNH_52076 | X | | Turbinates | Bone | 3.1 | 3.3 | 4.8 | 38.9% | 61.1% | 687 | 18 | 1107 | | | 64% | 73% | 73% |
| H_DRC_038 | AMNH_52077 | X | X | Tooth | Bone | 1.6 | 9.6 | 18.1 | 93.5% | 6.5% | 1628 | 66 | 118 | Hap_9 | W | 93% | 100% | 100% |
| H_DRC_039 | AMNH_52078 | X | X | Bone | Bone | 11.5 | 2.7 | 142.3 | 99.1% | 0.9% | 1700 | 96 | 16 | Hap_16 | W | 100% | 100% | 100% |
| H_DRC_040 | AMNH_52079 | X | X | Tooth | Bone | 2.4 | 2.2 | 297.7 | 99.6% | 0.4% | 1711 | 94 | 7 | Hap_12 | W | 93% | 100% | 100% |
| H_DRC_041 | AMNH_52080 | X | | Tooth | Bone | 2.7 | 2.8 | 5.0 | 42.4% | 57.6% | 740 | 29 | 1043 | | | 71% | 82% | 82% |
| H_DRC_042 | AMNH_52081 | X | X | Bone | Bone | 25.0 | 2.5 | 158.3 | 99.6% | 0.4% | 1710 | 95 | 7 | Hap_15 | W | 100% | 100% | 100% |
| H_DRC_043 | AMNH_52082 | X | X | Tooth | Bone | 7.8 | 3.5 | 326.9 | 99.7% | 0.3% | 1711 | 96 | 5 | Hap_14 | W | 79% | 91% | 91% |
| H_KEN_044 | AMNH_54370 | X | X | Bone | Bone | 28.0 | 2.0 | 100.8 | 99.4% | 0.6% | 1718 | 83 | 11 | Hap_29 | E | 100% | 100% | 100% |
| H_KEN_045 | AMNH_54371 | X | X | Bone | Bone | 11.0 | 2.2 | 111.5 | 99.1% | 0.9% | 1714 | 82 | 16 | Hap_30 | E | 100% | 100% | 100% |
| H_KEN_046 | AMNH_54372 | X | X | Bone | Bone | 10.4 | 3.3 | 17.6 | 96.0% | 4.0% | 1680 | 60 | 72 | Hap_24 | E | 100% | 100% | 100% |
| H_KEN_047 | AMNH_54393 | X | | Bone | Bone | 65.9 | 4.0 | 2.7 | 7.8% | 92.2% | 135 | 7 | 1670 | | | 93% | 100% | 100% |
| H_KEN_048 | AMNH_54394 | X | | Bone | Bone | 34.8 | 3.7 | 25.2 | 45.0% | 55.0% | 607 | 209 | 996 | | | 100% | 100% | 100% |
| H_KEN_049 | AMNH_54395 | X | | Bone | Bone | 67.1 | 2.9 | 7.9 | 30.7% | 69.3% | 401 | 155 | 1256 | | | 93% | 100% | 100% |
| H_GIR_050 | AMNH_54995 | X | X | Dried Tissue | Bone | 1.9 | 2.9 | 255.2 | 97.9% | 2.1% | 1685 | 89 | 38 | Hap_1 | W-N | 79% | 91% | 91% |
| H_GIR_051 | AMNH_54996 | X | X | Dried Tissue | Bone | 8.2 | 2.2 | 957.3 | 99.4% | 0.6% | 1704 | 98 | 10 | Hap_3 | W-N | 79% | 91% | 91% |
| H_CPT_052 | AMNH_6260 | | | Bone | Bone | 22.0 | 2.0 | 69.3 | 99.3% | 0.7% | 1724 | 76 | 12 | | | 100% | 100% | 100% |
| H_CPT_053 | AMNH_6282 | | | Turbinates | Bone | 16.5 | 2.0 | 16.9 | 91.5% | 8.5% | 1601 | 57 | 154 | | | 100% | 100% | 100% |
| H_GIR_054 | AMNH_63955 | X | X | Tooth | Bone | 8.1 | 2.6 | 231.9 | 99.4% | 0.6% | 1705 | 97 | 10 | Hap_2 | W-N | 100% | 100% | 100% |
| H_CPT_055 | AMNH_65 | | | Tooth | Bone | 12.5 | 1.8 | 49.0 | 99.3% | 0.7% | 1780 | 19 | 13 | | | 93% | 100% | 100% |
| H_CPT_056 | AMNH_70171 | | | Bone | Bone | 12.2 | 5.7 | 55.5 | 98.7% | 1.3% | 1697 | 91 | 24 | | | 86% | 91% | 91% |
| H_KEN_057 | AMNH_70347 | X | X | Bone | Bone | 17.7 | 2.4 | 93.1 | 99.4% | 0.6% | 1704 | 97 | 11 | Hap_18 | W | 93% | 100% | 100% |
| H_ANG_058 | AMNH_80609 | X | | Bone | Bone | 72.5 | 2.0 | 5.1 | 44.6% | 55.4% | 795 | 14 | 1003 | | | 100% | 100% | 100% |
| H_RSA_059 | AMNH_81836 | X | X | Tooth | Bone | 7.3 | 2.2 | 166.0 | 99.6% | 0.4% | 1720 | 84 | 8 | Hap_69 | S | 93% | 100% | 100% |
| H_RSA_060 | AMNH_81837 | X | X | Tooth | Bone | 5.3 | 2.4 | 400.0 | 99.7% | 0.3% | 1726 | 80 | 6 | Hap_68 | S | 93% | 91% | 91% |
| H_RSA_061 | AMNH_81839 | X | X | Tooth | Bone | 1.4 | 2.0 | 54.4 | 99.1% | 0.9% | 1718 | 77 | 17 | Hap_67 | S | 79% | 91% | 91% |
| H_RSA_062 | AMNH_81840 | X | X | Tooth | Bone | 283.0 | 2.5 | 305.2 | 99.7% | 0.3% | 1726 | 80 | 6 | Hap_68 | S | 100% | 100% | 100% |
| H_RSA_063 | AMNH_81841 | X | X | Tooth | Bone | 4.2 | 2.8 | 430.6 | 99.8% | 0.2% | 1788 | 20 | 4 | Hap_50 | M | 79% | 91% | 91% |
| H_RSA_064 | AMNH_81842 | X | X | Tooth | Bone | 8.0 | 2.1 | 1703.9 | 99.9% | 0.1% | 1790 | 20 | 2 | Hap_50 | M | 93% | 100% | 100% |
| H_RSA_065 | AMNH_81843 | X | X | Tooth | Bone | 18.1 | 1.9 | 202.9 | 99.7% | 0.3% | 1787 | 20 | 5 | Hap_50 | M | 100% | 100% | 100% |
| H_RSA_066 | AMNH_81844 | X | X | Bone | Bone | 6.4 | 2.7 | 161.1 | 99.1% | 0.9% | 1712 | 84 | 16 | Hap_70 | S | 93% | 100% | 100% |
| H_CAR_067 | AMNH_83410 | X | X | Tooth | Bone | 19.6 | 1.9 | 174.3 | 99.6% | 0.4% | 1709 | 96 | 7 | Hap_14 | W | 100% | 100% | 100% |
| H_CPT_068 | AMNH_8355 | | | Turbinates | Bone | 27.3 | 4.8 | 4.9 | 38.0% | 62.0% | 669 | 20 | 1123 | | | 86% | 100% | 100% |
| H_BOT_069 | AMNH_83617 | X | X | Turbinates | Bone | 3.5 | 3.6 | 15.7 | 95.8% | 4.2% | 1671 | 64 | 77 | Hap_73 | S | 93% | 100% | 100% |
| H_BOT_070 | AMNH_83618 | X | X | Turbinates | Bone | 8.0 | 2.0 | 73.9 | 99.1% | 0.9% | 1714 | 82 | 16 | Hap_88 | S | 100% | 100% | 100% |
| H_BOT_071 | AMNH_83619 | X | X | Tooth | Bone | 29.1 | 2.2 | 108.9 | 99.7% | 0.3% | 1726 | 80 | 6 | Hap_78 | S | 100% | 100% | 100% |
| H_BOT_072 | AMNH_83620 | X | X | Tooth | Bone | 1.4 | 1.9 | 17.0 | 90.1% | 9.9% | 1604 | 28 | 180 | Hap_35 | S | 43% | 55% | 55% |
| H_BOT_073 | AMNH_83621 | X | X | Turbinates | Bone | 15.4 | 1.9 | 51.3 | 98.8% | 1.2% | 1718 | 73 | 21 | Hap_80 | S | 100% | 100% | 100% |
| H_BOT_074 | AMNH_83622 | X | X | Bone | Bone | 8.7 | 2.2 | 81.5 | 99.0% | 1.0% | 1717 | 76 | 19 | Hap_81 | S | 100% | 100% | 100% |
| H_BOT_075 | AMNH_83623 | X | X | Bone | Bone | 6.9 | 2.3 | 74.8 | 99.1% | 0.9% | 1715 | 80 | 17 | Hap_87 | S | 100% | 100% | 100% |
| H_BOT_076 | AMNH_83624 | X | X | Turbinates | Bone | 20.9 | 2.0 | 56.4 | 98.8% | 1.2% | 1713 | 78 | 21 | Hap_85 | S | 100% | 100% | 100% |
| H_BOT_077 | AMNH_83625 | X | X | Tooth | Bone | 14.4 | 2.1 | 138.2 | 99.7% | 0.3% | 1724 | 83 | 5 | Hap_89 | S | 86% | 91% | 91% |
| H_CPT_078 | AMNH_8364 | | | Tooth | Bone | 17.9 | 5.1 | 10.7 | 81.4% | 18.6% | 1428 | 47 | 337 | | | 79% | 82% | 82% |
| H_TAN_079 | AMNH_85140 | X | X | Dried Tissue | Bone | 8.8 | 6.2 | 115.5 | 99.1% | 0.9% | 1774 | 21 | 17 | Hap_65 | M | 86% | 100% | 100% |
| H_TAN_080 | AMNH_85141-L | X | X | Tooth | Bone | 3.3 | 1.9 | 46.7 | 99.1% | 0.9% | 1774 | 22 | 16 | Hap_66 | M | 93% | 100% | 100% |
| H_TAN_081 | AMNH_85141-N | X | X | Bone | Bone | 12.8 | 2.2 | 26.7 | 96.4% | 3.6% | 1727 | 19 | 66 | Hap_56 | M | 100% | 100% | 100% |

### B.1.e: Historical Lions – Results (Page 2 of 2)

| | | Datasets | | Original | Extraction | NGS Coverage | | | mtDNA Sequence Variation | | | | | | | STR % Amplification | | |
|---|---|---|---|---|---|---|---|---|---|---|---|---|---|---|---|---|---|---|
| Project ID | Museum ID | nDNA | mtDNA | Material | Protocol | ng/uL | DNA | mtDNA | % Called | %Missing | REF | ALT | ? | Haplotype | Clade | 14 Loci | 8 Loci | 9 Loci |
| H_TAN_082 | AMNH_85142-L | X | | Tooth | Bone | 5.1 | 2.3 | 2.8 | 11.2% | 88.8% | 202 | 1 | 1609 | | | 71% | 82% | 82% |
| H_TAN_083 | AMNH_85142-N | X | X | Bone | Bone | 10.0 | 2.2 | 30.6 | 97.8% | 2.2% | 1751 | 21 | 40 | Hap_65 | M | 100% | 100% | 100% |
| H_TAN_084 | AMNH_85143 | X | X | Bone | Bone | 7.1 | 2.3 | 55.6 | 98.8% | 1.2% | 1769 | 22 | 21 | Hap_66 | M | 100% | 100% | 100% |
| H_TAN_085 | AMNH_85144 | X | X | Bone | Bone | 5.5 | 2.4 | 17.7 | 93.3% | 6.7% | 1674 | 16 | 122 | Hap_54 | M | 86% | 100% | 100% |
| H_TAN_086 | AMNH_85145 | X | X | Tooth | Bone | 1.8 | 1.8 | 22.1 | 98.3% | 1.7% | 1762 | 20 | 30 | Hap_60 | M | 86% | 91% | 91% |
| H_TAN_087 | AMNH_85146 | X | X | Bone | Bone | 142.0 | 1.9 | 94.9 | 99.4% | 0.6% | 1778 | 23 | 11 | Hap_59 | M | 100% | 100% | 100% |
| H_TAN_088 | AMNH_85147 | X | X | Dried Tissue | Bone | 9.5 | 2.1 | 327.9 | 99.6% | 0.4% | 1782 | 22 | 8 | Hap_66 | M | 57% | 64% | 64% |
| H_TAN_089 | AMNH_85148 | X | X | Bone | Bone | 6.6 | 2.9 | 159.1 | 98.8% | 1.2% | 1769 | 21 | 22 | Hap_64 | M | 100% | 100% | 100% |
| H_TAN_090 | AMNH_85149 | X | X | Bone | Bone | 9.3 | 2.2 | 24.6 | 94.6% | 5.4% | 1697 | 17 | 98 | Hap_45 | M | 100% | 100% | 100% |
| H_KEN_091 | AMNH_88632 | X | X | Tooth | Bone | 2.8 | 2.0 | 51.5 | 99.1% | 0.9% | 1717 | 78 | 17 | Hap_27 | E | 100% | 100% | 100% |
| H_KEN_092 | AMNH_88633 | X | X | Tooth | Bone | 2.3 | 1.8 | 29.9 | 97.5% | 2.5% | 1700 | 66 | 46 | Hap_25 | E | 64% | 82% | 82% |
| H_KEN_093 | AMNH_88634 | X | | Tooth | Bone | 9.2 | 1.7 | 7.8 | 75.6% | 24.4% | 1353 | 17 | 442 | | | 86% | 91% | 91% |
| H_KEN_094 | AMNH_88635 | X | X | Tooth | Bone | 4.4 | 2.6 | 222.5 | 99.3% | 0.7% | 1718 | 82 | 12 | Hap_31 | E | 93% | 100% | 100% |
| H_KEN_095 | AMNH_88636 | X | X | Tooth | Bone | 11.5 | 2.1 | 41.5 | 98.9% | 1.1% | 1711 | 81 | 20 | Hap_28 | E | 100% | 100% | 100% |
| H_KEN_096 | AMNH_88637 | X | X | Tooth | Bone | 3.8 | 2.1 | 64.0 | 99.3% | 0.7% | 1717 | 82 | 13 | Hap_26 | E | 100% | 100% | 100% |
| H_CPT_097 | CM_1461 | | | Tooth | Bone | 5.0 | 2.9 | 14.7 | 92.2% | 7.8% | 1598 | 72 | 142 | | | 64% | 82% | 82% |
| H_CPT_098 | CM_1564 | | | Dried Tissue | DNeasy | 3.5 | 2.4 | 91.1 | 98.3% | 1.7% | 1696 | 86 | 30 | | | 93% | 100% | 100% |
| H_CPT_099 | CM_1565 | | | Dried Tissue | DNeasy | 3.4 | 2.1 | 59.0 | 99.0% | 1.0% | 1703 | 90 | 19 | | | 86% | 91% | 91% |
| H_CPT_100 | CM_1825 | | | Bone | Bone | 10.5 | 5.0 | 17.4 | 94.3% | 5.7% | 1650 | 59 | 103 | | | 50% | 64% | 64% |
| H_UNK_101 | CM_184 | | | Tooth | Bone | 1.6 | 1.9 | 9.6 | 76.3% | 23.7% | 1334 | 48 | 430 | | | 43% | 55% | 55% |
| H_UNK_102 | CM_185 | | | Dried Tissue | DNeasy | 1.2 | 2.9 | 3.3 | 9.6% | 90.4% | 173 | 1 | 1638 | | | 36% | 45% | 45% |
| H_UNK_103 | CM_31 | | | Turbinates | Bone | 15.1 | 4.8 | 2.3 | 3.1% | 96.9% | 56 | 0 | 1756 | | | 93% | 100% | 100% |
| H_ETH_104 | CM_5868 | X | X | Tooth | Bone | 1.8 | 2.0 | 21.5 | 98.2% | 1.8% | 1759 | 20 | 33 | Hap_46 | M | 57% | 73% | 73% |
| H_TAN_105 | CM_5897 | X | X | Bone | Bone | 2.9 | 3.1 | 21.0 | 93.0% | 7.0% | 1668 | 18 | 126 | Hap_44 | M | 86% | 91% | 91% |
| H_TAN_106 | CM_5898 | X | X | Dried Tissue | DNeasy | 6.6 | 2.1 | 98.5 | 99.3% | 0.7% | 1779 | 21 | 12 | Hap_65 | M | 71% | 82% | 82% |
| H_TAN_107 | CM_5899 | X | X | Turbinates | Bone | 19.8 | 2.1 | 18.7 | 95.8% | 4.2% | 1715 | 20 | 77 | Hap_58 | M | 100% | 100% | 100% |
| H_TAN_108 | FMNH_127836 | X | X | Dried Tissue | DNeasy | 2.6 | 2.9 | 21.8 | 94.7% | 5.3% | 1698 | 18 | 96 | Hap_53 | M | 64% | 82% | 82% |
| H_TAN_109 | FMNH_127837 | X | | Dried Tissue | DNeasy | 2.5 | 5.1 | 4.1 | 17.0% | 83.0% | 306 | 2 | 1504 | | | 50% | 55% | 55% |
| H_TAN_110 | FMNH_127838 | X | | Dried Tissue | DNeasy | 8.4 | 5.9 | 3.4 | 12.1% | 87.9% | 216 | 3 | 1593 | | | 71% | 91% | 91% |
| H_TAN_111 | FMNH_127839 | X | X | Dried Tissue | DNeasy | 5.3 | 2.9 | 59.4 | 98.0% | 2.0% | 1755 | 20 | 37 | Hap_62 | M | 43% | 55% | 55% |
| H_TAN_112 | FMNH_127840 | X | | Dried Tissue | DNeasy | 2.1 | 8.8 | 2.7 | 1.2% | 98.8% | 22 | 0 | 1790 | | | 36% | 45% | 45% |
| H_SOM_113 | FMNH_1443 | X | X | Dried Tissue | DNeasy | 1.3 | 3.1 | 37.2 | 94.3% | 5.7% | 1627 | 81 | 104 | Hap_10 | W | 57% | 73% | 73% |
| H_KEN_114 | FMNH_20756 | X | | Dried Tissue | DNeasy | 1.3 | 2.8 | 7.0 | 5.2% | 94.8% | 66 | 29 | 1717 | | | 64% | 73% | 73% |
| H_KEN_115 | FMNH_20757 | X | X | Dried Tissue | DNeasy | 2.5 | 2.4 | 57.3 | 96.9% | 3.1% | 1734 | 21 | 57 | Hap_63 | M | 100% | 100% | 100% |
| H_KEN_116 | FMNH_20758 | X | X | Dried Tissue | DNeasy | 1.2 | 2.9 | 34.9 | 97.7% | 2.3% | 1751 | 20 | 41 | Hap_62 | M | 57% | 73% | 73% |
| H_KEN_117 | FMNH_20760 | X | X | Dried Tissue | DNeasy | 1.5 | 2.4 | 312.0 | 99.7% | 0.3% | 1784 | 22 | 6 | Hap_66 | M | 71% | 82% | 82% |
| H_KEN_118 | FMNH_20762 | X | X | Dried Tissue | DNeasy | 1.5 | 2.8 | 97.4 | 98.4% | 1.6% | 1762 | 21 | 29 | Hap_61 | M | 86% | 100% | 100% |
| H_SDN_119 | FMNH_30778 | X | | Dried Tissue | DNeasy | 1.0 | 2.4 | 2.1 | 1.2% | 98.8% | 21 | 0 | 1791 | | | 64% | 73% | 73% |
| H_GIR_120 | FMNH_31121 | X | X | Dried Tissue | DNeasy | 2.1 | 2.6 | 287.3 | 99.4% | 0.6% | 1703 | 98 | 11 | Hap_4 | W-N | 100% | 100% | 100% |
| H_TAN_121 | FMNH_33479 | X | | Dried Tissue | Bone | 3.7 | 3.0 | 12.6 | 85.8% | 14.2% | 1544 | 10 | 258 | | | 7% | 9% | 9% |
| H_TAN_122 | FMNH_33480 | X | | Dried Tissue | DNeasy | 3.6 | 3.8 | 5.2 | 45.8% | 54.2% | 820 | 9 | 983 | | | 71% | 91% | 91% |
| H_TAN_123 | FMNH_35131 | X | X | Dried Tissue | DNeasy | 1.9 | 3.6 | 182.5 | 99.6% | 0.4% | 1783 | 22 | 7 | Hap_66 | M | 50% | 64% | 64% |
| H_TAN_124 | FMNH_35132 | X | X | Dried Tissue | DNeasy | 6.5 | 3.5 | 1459.0 | 99.7% | 0.3% | 1784 | 22 | 6 | Hap_66 | M | 29% | 36% | 36% |
| H_TAN_125 | FMNH_35133 | X | X | Dried Tissue | DNeasy | 12.1 | 1.8 | 140.0 | 99.7% | 0.3% | 1784 | 22 | 6 | Hap_66 | M | 100% | 100% | 100% |
| H_TAN_126 | FMNH_35134 | X | X | Dried Tissue | DNeasy | 6.2 | 2.2 | 562.5 | 99.9% | 0.1% | 1788 | 22 | 2 | Hap_66 | M | 100% | 100% | 100% |
| H_BOT_127 | FMNH_35739 | X | X | Dried Tissue | DNeasy | 1.4 | 3.3 | 204.2 | 99.2% | 0.8% | 1777 | 20 | 15 | Hap_50 | M | 86% | 91% | 91% |
| H_BOT_128 | FMNH_35740 | X | | Dried Tissue | DNeasy | 2.3 | 7.0 | 4.8 | 34.0% | 66.0% | 616 | 0 | 1196 | | | 57% | 64% | 64% |
| H_BOT_129 | FMNH_35741 | X | X | Dried Tissue | DNeasy | 4.3 | 2.3 | 239.7 | 99.7% | 0.3% | 1727 | 79 | 6 | Hap_79 | S | 86% | 100% | 100% |
| H_BOT_130 | FMNH_35742 | X | | Dried Tissue | DNeasy | 6.8 | 2.2 | 10.9 | 81.5% | 18.5% | 1430 | 47 | 335 | | | 71% | 91% | 91% |
| H_BOT_131 | FMNH_35743 | X | X | Dried Tissue | DNeasy | 3.2 | 3.1 | 369.7 | 99.2% | 0.8% | 1718 | 79 | 15 | Hap_79 | S | 57% | 64% | 64% |
| H_RSA_132 | FMNH_38134 | X | X | Dried Tissue | DNeasy | 12.0 | 2.1 | 82.4 | 99.3% | 0.7% | 1723 | 76 | 13 | Hap_75 | S | 100% | 100% | 100% |
| H_BOT_133 | FMNH_41405 | X | X | Dried Tissue | DNeasy | 4.5 | 2.5 | 262.1 | 99.6% | 0.4% | 1725 | 80 | 7 | Hap_82 | S | 100% | 100% | 100% |
| H_MLI_134 | FMNH_42129 | X | | Dried Tissue | DNeasy | 1.7 | 2.5 | 11.2 | 74.5% | 25.5% | 1321 | 29 | 462 | | | 71% | 82% | 82% |
| H_KEN_135 | FMNH_75608 | X | | Dried Tissue | DNeasy | 28.4 | 5.2 | 1.9 | 0.6% | 99.4% | 10 | 0 | 1802 | | | 21% | 27% | 27% |
| H_KEN_136 | FMNH_75609 | X | | Dried Tissue | DNeasy | 5.3 | 8.5 | 9.5 | 81.6% | 18.4% | 1466 | 12 | 334 | | | 50% | 55% | 55% |
| H_BOT_137 | FMNH_89926 | X | X | Dried Tissue | DNeasy | 3.3 | 2.7 | 231.3 | 99.6% | 0.4% | 1724 | 81 | 7 | Hap_84 | S | 100% | 100% | 100% |
| H_SDN_138 | KBIN_469459 | X | | Skull | *** | | | | | | | | | | | 14% | 18% | 18% |
| H_ZAM_139 | KBIN_504512 | X | | Skull | *** | | | | | | | | | | | 86% | 91% | 91% |
| H_TAN_140 | KU_105216 | X | X | Bone | Bone | 1.0 | 1.9 | 62.4 | 98.3% | 1.7% | 1767 | 15 | 30 | Hap_52 | M | 79% | 82% | 82% |
| H_TAN_141 | KU_105217 | X | X | Bone | Bone | 125.0 | 2.6 | 125.6 | 99.4% | 0.6% | 1781 | 21 | 10 | Hap_65 | M | 100% | 100% | 100% |
| H_KEN_142 | LACM_51295 | X | X | Bone | Bone | 97.1 | 2.1 | 11.8 | 88.1% | 11.9% | 1578 | 18 | 216 | Hap_55 | M | 93% | 100% | 100% |
| H_KEN_143 | LACM_51296 | X | | Bone | Bone | 12.6 | 2.1 | 9.5 | 80.9% | 19.1% | 1452 | 14 | 346 | | | 100% | 100% | 100% |
| H_KEN_144 | LACM_51297 | X | | Bone | DNeasy | 1.3 | 3.5 | 2.1 | 1.2% | 98.8% | 21 | 0 | 1791 | | | 36% | 36% | 36% |
| H_TAN_145 | MVZ_96804 | X | X | Turbinates | Bone | 3.2 | 3.8 | 14.7 | 95.6% | 4.4% | 1711 | 21 | 80 | Hap_48 | M | 64% | 73% | 73% |
| H_ANG_146 | RMNH_45281 | X | | Skull | *** | | | | | | | | | | | 71% | 82% | 82% |
| H_NRT_147 | RMNH_45282 | X | | Hide | *** | | | | | | | | | | | 50% | 55% | 55% |
| H_MOZ_148 | RMNH_45284 | X | | Skull | *** | | | | | | | | | | | 36% | 45% | 45% |
| H_MOZ_149 | RMNH_45285 | X | | Skull | *** | | | | | | | | | | | 57% | 64% | 64% |
| H_NAM_150 | S_581971 | X | | Skull | *** | | | | | | | | | | | 93% | 100% | 100% |
| H_NRT_151 | S_585287 | X | | Skull | *** | | | | | | | | | | | 43% | 45% | 45% |
| H_DRC_152 | S_595059 | X | | Skull | *** | | | | | | | | | | | 64% | 73% | 73% |
| H_TAN_153 | YPM_2057 | X | X | Hide | Bone | 1.3 | 2.7 | 228.0 | 97.6% | 2.4% | 1748 | 21 | 43 | Hap_57 | M | 71% | 82% | 82% |
| H_KEN_154 | YPM_3189 | X | X | Hide | Bone | 4.5 | 2.5 | 372.5 | 99.2% | 0.8% | 1775 | 22 | 15 | Hap_66 | M | 57% | 64% | 64% |
| H_KEN_155 | YPM_3217 | X | X | Hide | Bone | 10.6 | 2.3 | 585.0 | 99.4% | 0.6% | 1719 | 82 | 11 | Hap_33 | E | 93% | 100% | 100% |
| H_KEN_156 | YPM_3218 | X | X | Hide | Bone | 4.4 | 2.2 | 185.1 | 99.1% | 0.9% | 1778 | 17 | 17 | Hap_36 | M | 93% | 100% | 100% |
| H_KEN_157 | YPM_5251 | X | X | Dried Tissue | Bone | | 2.3 | 321.9 | 99.4% | 0.6% | 1720 | 81 | 11 | Hap_34 | E | 79% | 91% | 91% |
| H_CPT_158 | YPM_6943 | | | Bone | Bone | 19.7 | 2.5 | 30.4 | 92.9% | 7.1% | 1615 | 69 | 128 | | | 100% | 100% | 100% |
| H_UNK_159 | YPM_6952 | | | Bone | Bone | 18.9 | 2.5 | 3.0 | 10.7% | 89.3% | 191 | 2 | 1619 | | | 86% | 91% | 91% |
| H_KEN_160 | YPM_9570 | X | | Dried Tissue | DNeasy | 14.3 | 1.9 | 11.5 | 89.1% | 10.9% | 1601 | 13 | 198 | | | 100% | 100% | 100% |
| H_RSA_161 | ZMA_11753 | X | | Skull | *** | | | | | | | | | | | 50% | 55% | 55% |
| H_CAR_162 | ZMA_996 | X | | Skull | *** | | | | | | | | | | | 86% | 91% | 91% |

**S2. Step-by-step Hierarchical STRUCTURE:**

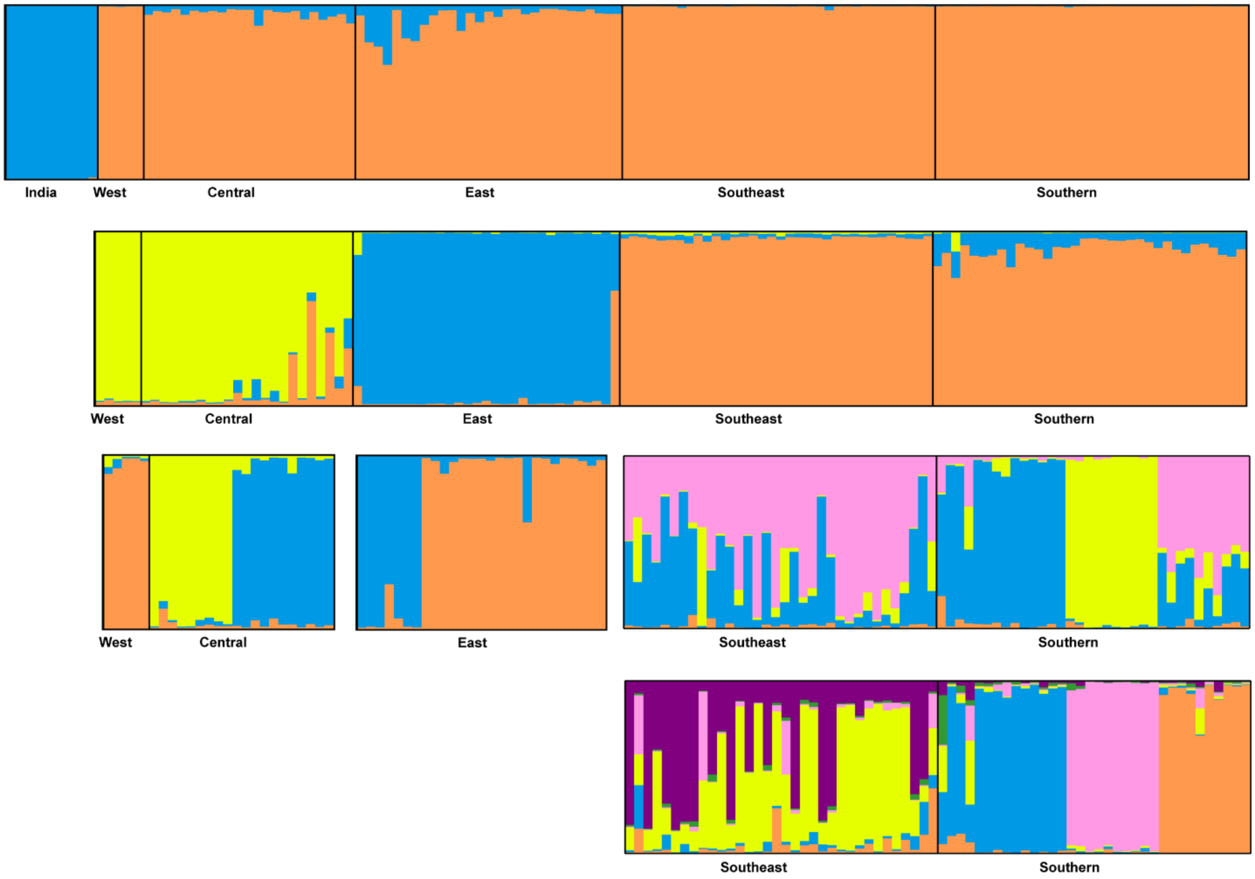

**S3. MD Diversity Statistics:** Nuclear genetic diversity for lion subpopulations defined during hierarchical structure analysis of the modern population. Color is based on when the population emerges during hierarchical structure analysis (see Figure.Structure). Dark gray encompasses all data. Light gray is the continental tier. Yellow is the subcontinental tier. Blue is the regional tier. Green in the local tier.

| | N | P (%) | A (AP) | Range | H | $H_O$ | $H_E$ | M |
|---|---|---|---|---|---|---|---|---|
| **All Data** | **135** | **100** | **11.6** | **30.0** | **0.80** | **0.64** | **0.80** | **0.32** |
| Asia | 10 | 22 | 1.3 (2.5) | 6.0 | 0.09 | 0.36 | 0.42 | 0.05 |
| Africa | 125 | 100 | 11.6 | 30.0 | 0.79 | 0.69 | 0.79 | 0.39 |
| Western | 28 | 100 | 5.6 | 19.5 | 0.74 | 0.61 | 0.74 | 0.18 |
| WES | 5 | 33 | 2.3 (3.0) | 18.7 | 0.53 | 0.93 | 0.79 | 0.09 |
| MID | 9 | 67 | 2.0 (2.5) | 13.7 | 0.46 | 0.76 | 0.69 | 0.07 |
| CEN | 14 | 44 | 3.6 (8.0) | 24.5 | 0.36 | 0.73 | 0.82 | 0.13 |
| Eastern | 29 | 100 | 6.1 | 20.9 | 0.66 | 0.62 | 0.66 | 0.22 |
| KEN | 7 | 33 | 1.3 (3.7) | 16.7 | 0.22 | 0.71 | 0.66 | 0.05 |
| TAN | 22 | 100 | 6.1 | 20.9 | 0.66 | 0.63 | 0.66 | 0.22 |
| Southern | 68 | 100 | 9.7 | 28.0 | 0.79 | 0.71 | 0.79 | 0.36 |
| Southeast | 34 | 100 | 7.7 | 26.4 | 0.76 | 0.71 | 0.76 | 0.26 |
| ZAE | 17 | 100 | 6.1 | 25.1 | 0.67 | 0.65 | 0.67 | 0.21 |
| ZAW | 17 | 100 | 6.1 | 22.4 | 0.77 | 0.76 | 0.77 | 0.22 |
| South | 14 | 100 | 6.9 | 24.0 | 0.78 | 0.80 | 0.78 | 0.24 |
| Southwest | 20 | 100 | 5.3 | 22.9 | 0.72 | 0.66 | 0.72 | 0.19 |
| KAL | 10 | 100 | 4.1 | 16.4 | 0.69 | 0.69 | 0.69 | 0.15 |
| ETO | 10 | 100 | 3.8 | 18.2 | 0.62 | 0.63 | 0.62 | 0.14 |

sample size (N)
percentage of polymorphic loci (P)
mean number of alleles per locus (A)
mean number of alleles per polymorphic locus if different than A (AP)
allelic range (Range)
genetic diversity of mean expected heterozygosity per locus (H)
mean observed heterozygosity per polymorphic locus ($H_O$)
mean expected heterozygosity per polymorphic locus ($H_E$)
Garza-Williamson modified index (M)

**S4. Additional HD PCoA Analyses**

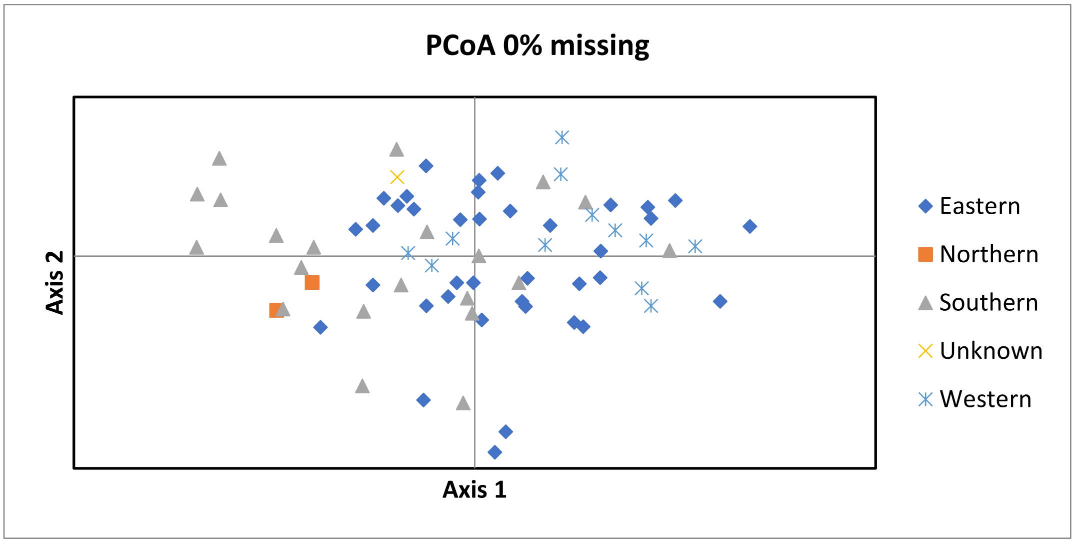


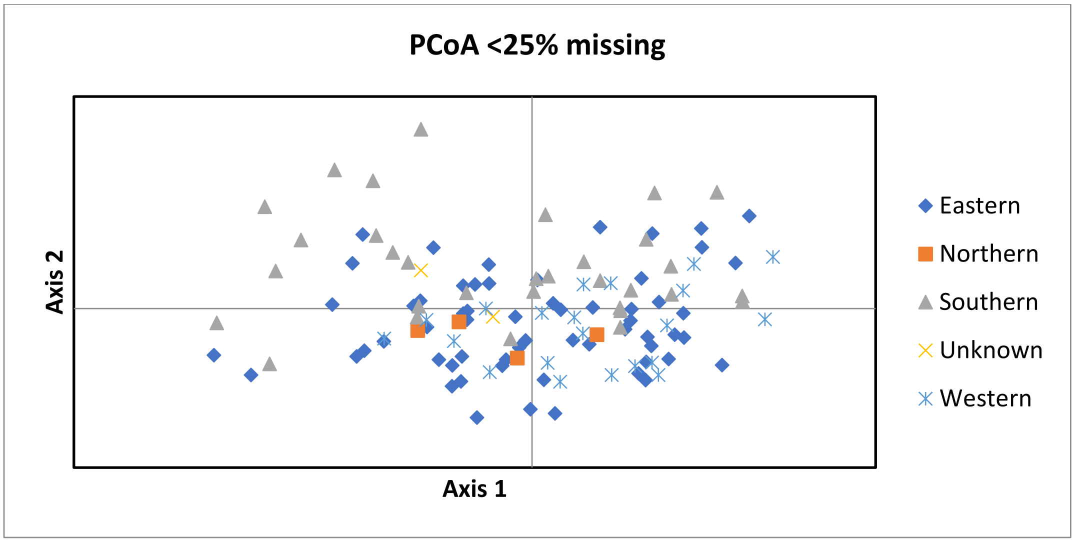


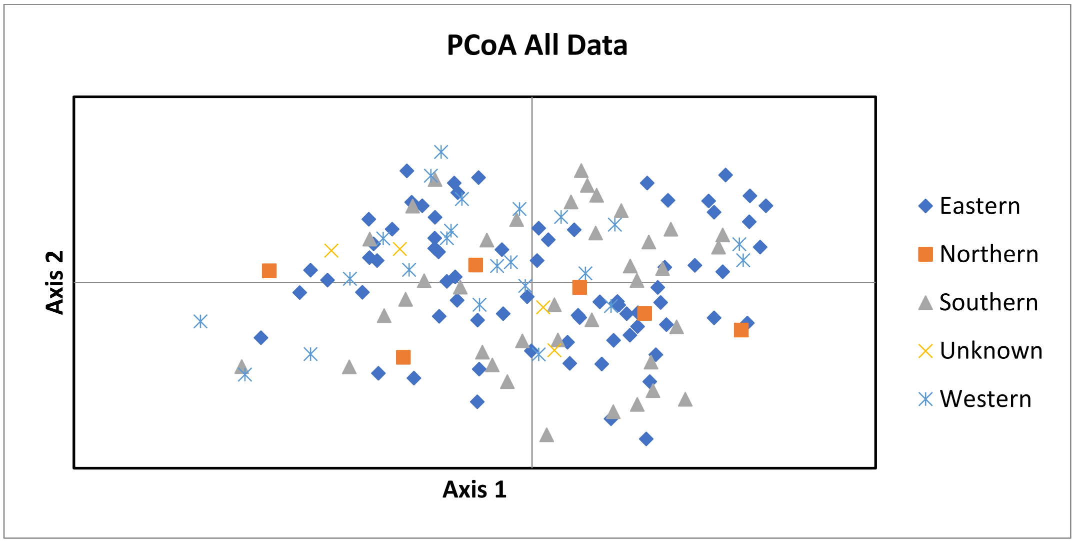

**S5.a. Mantel Test for Isolation-by-Distance of HD**

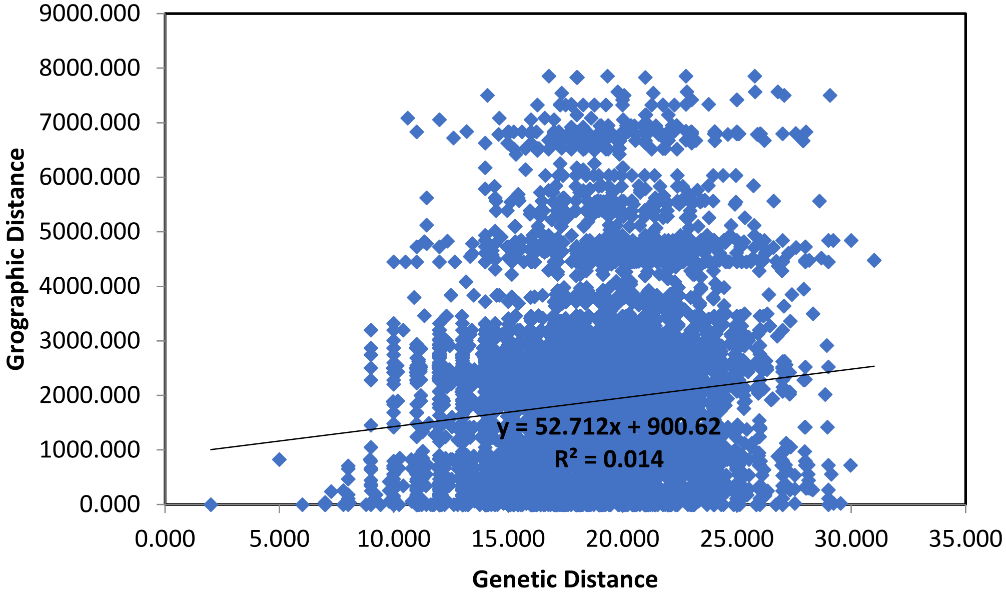


*Genetic distance tested against geographic distance (p-value = 0.060) in GenAlEx 6.503.*

**S5.b. Mantel Test for Isolation-by-Distance of MD**

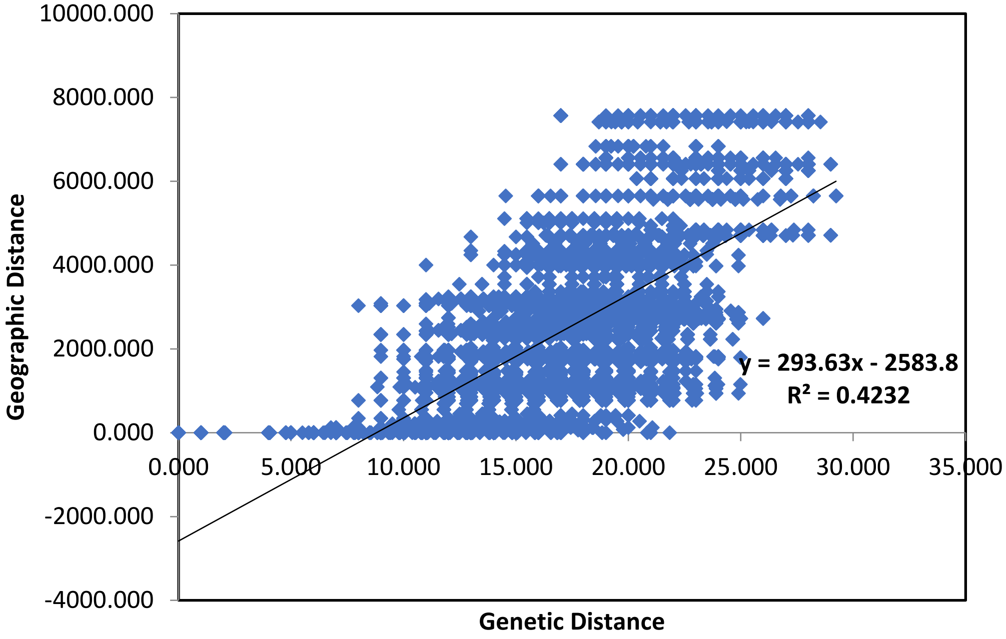


*Genetic distance tested against geographic distance (p-value = 0.010) in GenAlEx 6.503.*

**S6. Mitogenome PCA.** Principal Component Analysis of 280 variable sites from 121 lion mitogenomes. PC1=17%, PC2=12% and PC3=10% of the total variation. Color corresponds to conventionally recognized regions (See Figure 1).

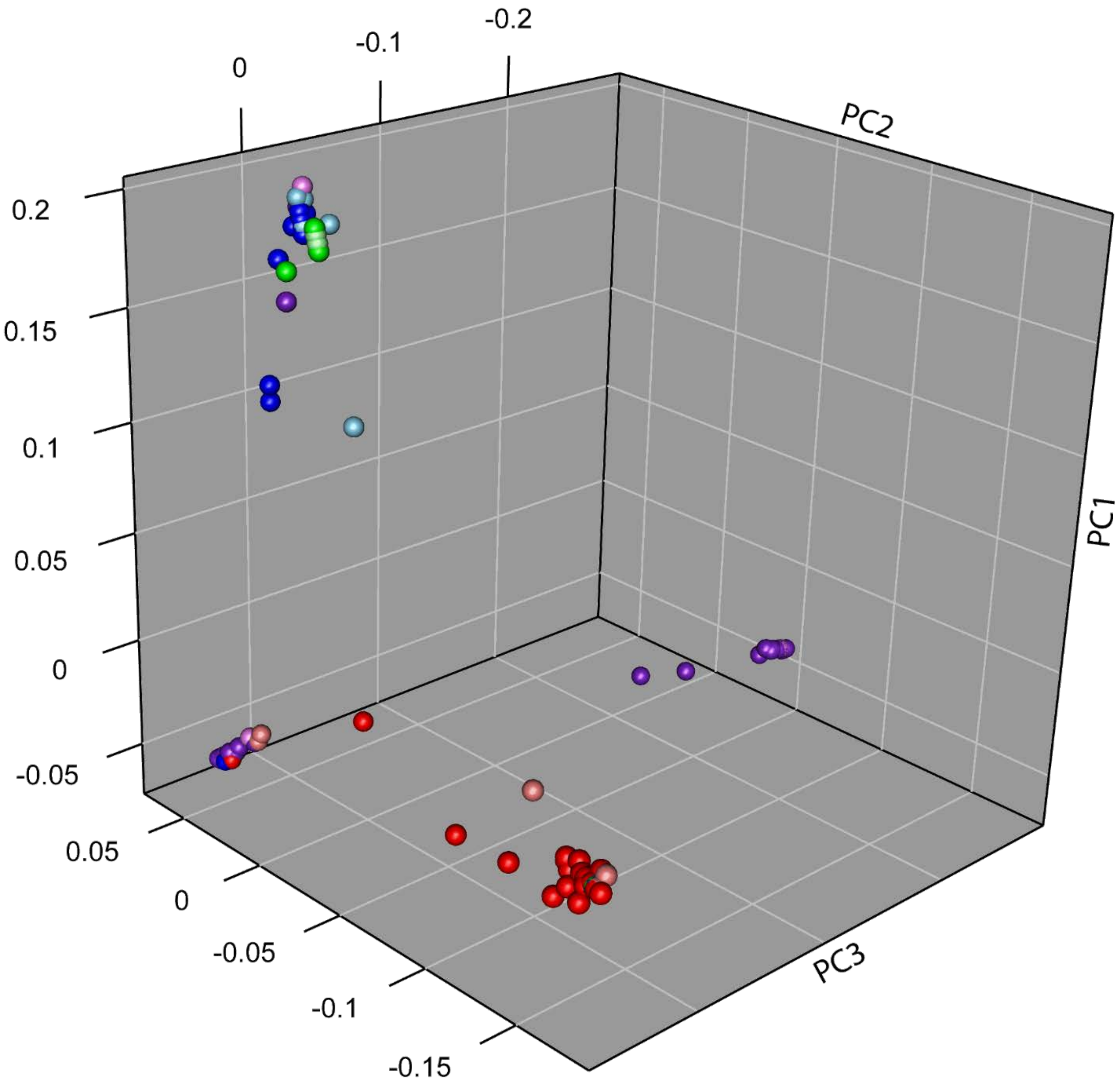

**S7. Mitogenome Haplotype Network.** Neighbor-joining haplotype network based on 280 variable sites from 121 lion mitogenomes. Color corresponds to conventionally recognized regions.

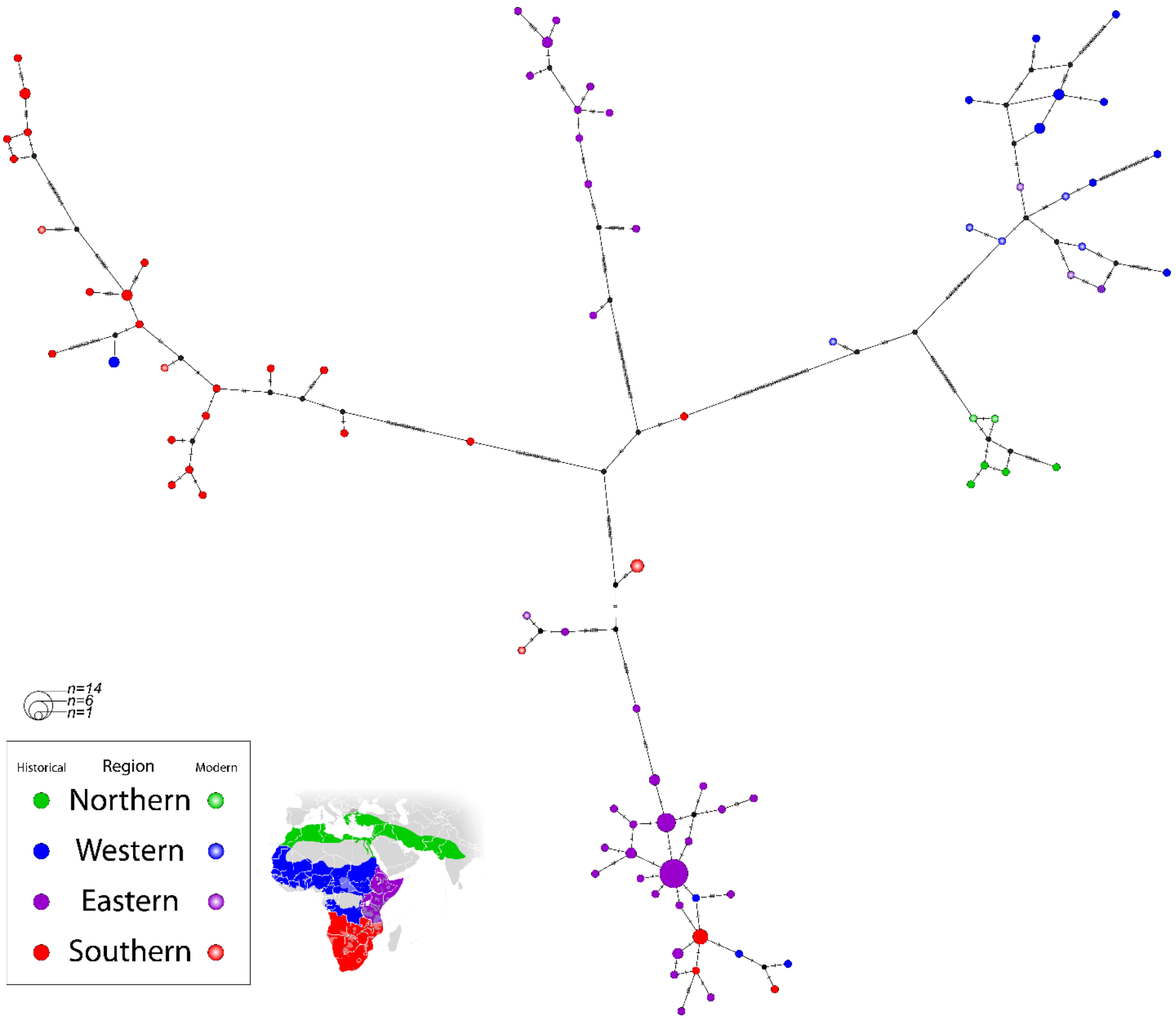

**S8. Comparison of mitochondrial population structure** in lions with previously published haplotype networks illustrating maintenance of mitochondrial structure.

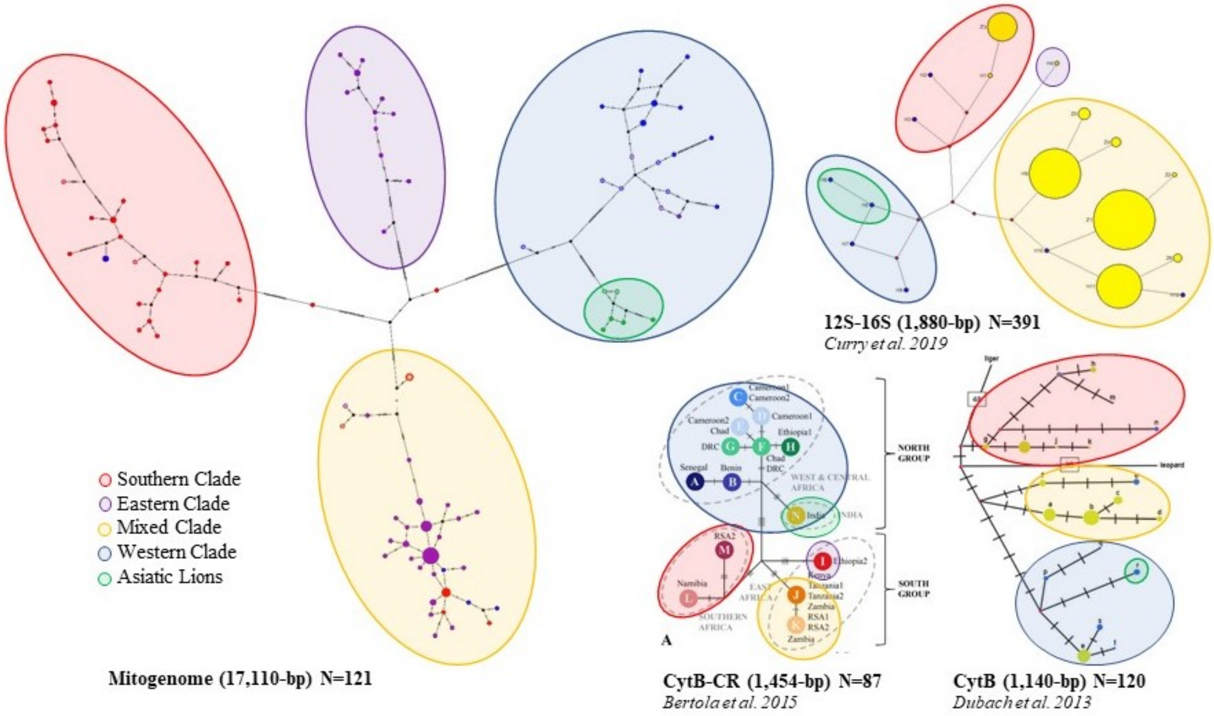

**S9. Table of Museums**: List of natural history museums that provided lion samples for the historical population inferences.

| Museum | *Location* | Provided | nDNA | mtDNA |
|---|---|---|---|---|
| *American Museum of Natural History (AMNH) | *New York, NY, USA* | 96 | 80 | 71 |
| *Carnegie Museum of Natural History (CM) | *Pittsburgh, PA, USA* | 11 | 4 | 4 |
| *Field Museum of Natural History (FMNH) | *Chicago, IL, USA* | 30 | 30 | 18 |
| *Kansas University Natural History Museum (KU) | *Lawrence, KS, USA* | 2 | 2 | 2 |
| *Natural History Museum of Los Angeles County (LACM) | *Los Angeles, CA, USA* | 3 | 3 | 1 |
| **Naturalis Biodiversity Center (RMNH) | *Leiden, Netherlands* | 4 | 4 | 0 |
| **Royal Belgian Institute of Natural Sciences (KBIN) | *Bruxelles, Belgium* | 2 | 2 | 0 |
| **Swedish Royal Museum of Natural History (S) | *Stockholm, Sweden* | 3 | 3 | 0 |
| *The Museum of Vertebrate Zoology at Berkeley (MVZ) | *Berkeley, CA, USA* | 1 | 1 | 1 |
| *Yale Peabody Museum (YPM) | *New Haven, CT, USA* | 8 | 6 | 5 |
| **Zoological Museum Amsterdam (ZMA) | *Amsterdam, Netherlands* | 2 | 2 | 0 |

*Collected by and DNA extraction done by Texas A&M University.

** Collected by and DNA extraction done by Leiden University (see Bertola *et al.* 2016).

**S10. Methods**

Nuclear DNA

***Historical Lion Dataset (HD) Amplification & Allele Calling***

Sample preparation and DNA extraction method was dependent on sample type. Skeletal remains followed protocols established by Curry and Derr 2019 (1). Dried tissue was washed with dd$H_2O$ and ethanol then allowed to dry completely. DNA was then extracted using the DNeasy Blood and Tissue Kit (Qiagen) following manufacturer's instructions for purification of total DNA from animal tissues.

Samples were all processed one at a time in a clean workstation (C.B.S. Scientific Model #P-048-202) with dedicated equipment for work with lion samples. Between each sample, the workstation and all reusable equipment were cleaned with dd$H_2O$, 10% bleach, and/or 95% ethanol then UV irradiated for 10-20 minutes. Consumables were all UV irradiated before use.

Historical lions were amplified at 14 Leo STRs developed by Curry and Derr 2019 (1) for use with historical DNA (Leo006, 008, 031, 045, 077, 085, 098, 126, 224, 230, 247, 281, 391, and 506). Loci with great than 75% amplification success across samples that also appear in the modern dataset (Leo006, 008, 085, 098, 126, 224, 230, 247, and 281) were used for downstream analysis.

***Modern Lion Dataset (MD) Allele Call Calibration***

There is sampling overlap between MD-1 (2), MD-2 (3), and MD-3 (4). Lions in MD-2 were included in MD-1 (2), though MD-2 has more loci than MD-1. Nine lions from the eastern subpopulation in Zambia were used in both MD-1 and MD-3. MD-3 includes an additional 8 lions from the Zambian eastern subpopulation and 17 lions from the Zambian western subpopulations chosen at random to represent the known subpopulations in Zambia (4) at comparable sample sizes to subpopulations sampled in MD-1 and MD-2.

STRs were necessary for increased amplification of the historical samples (5), therefore, MD-1 data were calibrated to MD-3. Two samples from MD-1 (M_CAM_0019 and M_BEN_0013) were amplified and genotyped using FCA microsatellite primers and STR primers for loci 006, 008, 031, 045, 077, 085, 098, 126, 224, 230, 247, 281, 391, and 506. The FCA, Leo, and published allele calls were compared to determine a correction for calibration of the allele calls for the two studies (Table 1). This correction was then applied to all other allele calls for the appropriate loci. The 9 lions that appear in both MD-1 and MD-3 provide a secondary check for the accuracy of the calibration.

**Table 1: STR Calibration Results**

| | | 006 | | 008 | | 031 | | 045 | | 077 | | 085 | | 098 | | 126 | | 224 | | 230 | | 247 | | 281 | | 391 | | 506 | |
|---|---|---|---|---|---|---|---|---|---|---|---|---|---|---|---|---|---|---|---|---|---|---|---|---|---|---|---|---|---|
| BEN | Leo | 122 | 136 | 113 | 131 | 198 | 198 | 80 | 96 | 106 | 106 | 86 | 92 | 102 | 102 | 125 | 127 | 92 | 94 | 86 | 86 | 114 | 126 | 223 | 223 | 178 | 178 | 171 | 191 |
| BEN | Predicted | 204 | 218 | 126 | 144 | | | 132 | 148 | 148 | 148 | 129 | 135 | 104 | 104 | 137 | 139 | 166 | 168 | 92 | 92 | 135 | 147 | 242 | 242 | | | | |
| BEN | FCA | | | 126 | 144 | | | 132 | 148 | 148 | 148 | 129 | 135 | 104 | 104 | 137 | 139 | 168 | 170 | | | 135 | 147 | 243 | 243 | | | | |
| BEN | FCA | | | 126 | 144 | | | 132 | 148 | 148 | 148 | 129 | 135 | 104 | 104 | 137 | 139 | 168 | 170 | | | 135 | 147 | 243 | 243 | | | | |
| BEN | Study | | | | | | | | | | | 145 | 151 | | | 155 | 157 | 184 | 186 | | | 160 | 172 | | | | | | |
| BEN | Leo Diff | | | | | | | | | | | 59 | 59 | | | 30 | 30 | 92 | 92 | | | 46 | 46 | | | | | | |
| BEN | FCA Diff | | | | | | | | | | | 16 | 16 | | | 18 | 18 | 16 | 16 | | | 25 | 25 | | | | | | |
| CAM | Leo | 110 | 136 | 111 | 111 | 194 | 198 | 80 | 100 | 104 | 112 | 80 | 80 | 92 | 102 | 137 | 139 | 78 | 78 | 86 | 86 | 126 | 128 | 225 | 225 | 182 | 182 | 187 | 187 |
| CAM | Predicted | 192 | 218 | 124 | 124 | | | 132 | 152 | 146 | 154 | 123 | 123 | 94 | 104 | 149 | 151 | 152 | 152 | 92 | 92 | 147 | 149 | 244 | 244 | | | | |
| CAM | FCA | | | 122 | 122 | | | 132 | 154 | 146 | 154 | 123 | 123 | 96 | 106 | 148 | 150 | 150 | 150 | 92 | 92 | 147 | 149 | 145 | 145 | | | | |
| CAM | FCA | | | 122 | 122 | | | 132 | 154 | 146 | 154 | 123 | 123 | 96 | 104 | 148 | 150 | 152 | 152 | 92 | 92 | 147 | 149 | 145 | 145 | | | | |
| CAM | Study | | | | | | | | | | | 141 | 141 | | | 167 | 169 | 170 | 170 | | | 172 | 174 | | | | | | |
| CAM | Leo Diff | | | | | | | | | | | 61 | 61 | | | 30 | 30 | 92 | 92 | | | 46 | 46 | | | | | | |
| CAM | FCA Diff | | | | | | | | | | | 18 | 18 | | | 19 | 19 | 20 | 20 | | | 25 | 25 | | | | | | |
| 254 | Leo | 94 | 104 | 113 | 131 | 192 | 196 | 80 | 80 | 98 | 102 | 80 | 94 | 102 | 102 | 125 | 129 | 86 | 86 | 80 | 88 | 122 | 122 | 211 | 229 | 194 | 198 | 193 | 197 |
| 254 | FCA | 176 | 186 | 126 | 144 | | | 132 | 132 | 140 | 144 | 123 | 137 | 104 | 104 | 137 | 141 | 160 | 160 | 86 | 94 | 143 | 143 | 230 | 248 | | | | |
| 254 | Diff | 82 | 82 | 13 | 13 | | | 52 | 52 | 42 | 42 | 43 | 43 | 2 | 2 | 12 | 12 | 74 | 74 | 6 | 6 | 21 | 21 | 19 | 19 | | | | |
| 387 | Leo | 104 | 104 | 127 | 127 | 196 | 196 | 80 | 80 | 102 | 104 | 80 | 80 | 104 | 108 | 131 | 143 | 78 | 86 | 78 | 82 | 114 | 116 | 213 | 217 | 174 | 190 | 191 | 193 |
| 387 | FCA | 186 | 186 | 140 | 140 | | | 132 | 132 | 144 | 146 | 123 | 123 | 106 | 110 | 143 | 155 | 152 | 160 | 84 | 88 | 135 | 137 | 232 | 236 | | | | |
| 387 | Diff | 82 | 82 | 13 | 13 | | | 52 | 52 | 42 | 42 | 43 | 43 | 2 | 2 | 12 | 12 | 74 | 74 | 6 | 6 | 21 | 21 | 19 | 19 | | | | |
| | Diff | 82 | 82 | 13 | 13 | | | 52 | 52 | 42 | 42 | 43 | 43 | 2 | 2 | 12 | 12 | 74 | 74 | 6 | 6 | 21 | 21 | 19 | 19 | | | | |
| Driscol | MCA | 192 | 192 | 126 | 126 | | | 133 | 133 | 150 | 150 | 124 | 124 | 111 | 111 | 140 | 140 | 173 | 173 | 101 | 101 | 140 | 140 | 123 | 123 | | | | |
| Driscol | Leo MCA | 124 | 124 | 111 | 111 | 196 | 196 | 80 | 80 | 104 | 104 | 80 | 80 | 102 | 102 | 125 | 125 | 86 | 86 | 88 | 88 | 128 | 128 | 213 | 213 | 174 | 174 | 193 | 193 |
| Driscol | Diff | 68 | 68 | 15 | 15 | | | 53 | 53 | 46 | 46 | 44 | 44 | 9 | 9 | 15 | 15 | 87 | 87 | 13 | 13 | 12 | 12 | -90 | -90 | | | | |
| | Cal Used | 84 | | 13 | | | | 53 | | 44 | | 44 | | 7 | | 13 | | 77 | | 13 | | 26 | | -90 | | | | | |

Mitochondrial DNA

***Whole Genome Sequencing***

To obtain mitogenomes, 155 samples were whole genome sequenced. Library prep and sequencing were performed by Texas A&M AgriLife Genomics and Bioinformatics Services. Library prep was done using NEBNext® Ultra™ II, designed for low input amounts, and sample libraries were sequenced on the Illumina HiSeq 4000. Sequence cluster identification, quality prefiltering, base calling, and uncertainty assessment were done in real time using Illumina's HCS 2.2.68 and RTA 1.18.66.3 software with default parameter settings.

Reads were mapped by SpeedSeq (6) to the lion genome reference (provided by Ellie Armstrong at Stanford University) with a mitogenome sequence (GenBank Accession KP001505.1) and representative nuclear mitochondrial sequence (personal communication, Dr. Laura Bertola) added as additional scaffolds. The Panthera lineage contains multiple nuclear mitochondrial DNA segments (numts) (7, 8). Despite divergence between the mitogenome and numt, mapping only short reads causes an overlap of reads covering approximately 6500-bps of the mitogenome (Figure 1). To correct for the presence of numts, a consensus of the reads that deviate from the mitogenome reference, created by Dr. Laura Bertola, was added to the reference genome as a representative numt scaffold.

**Figure 1: Mitogenome Alignments with and without NUMT Correction**

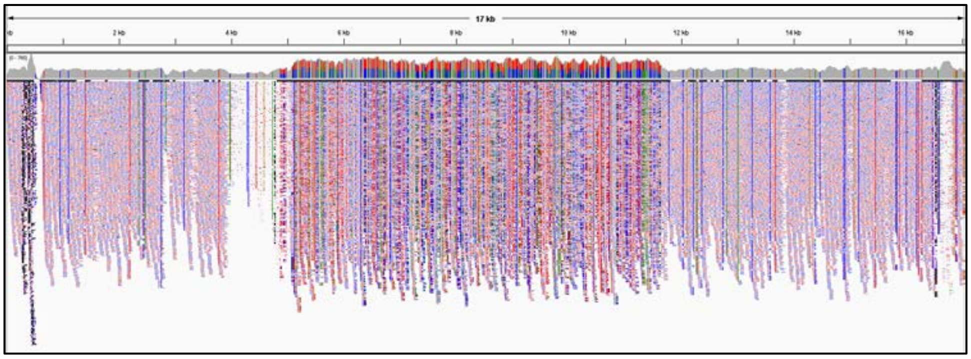

Historical Lion AMNH_81840 without numt correction

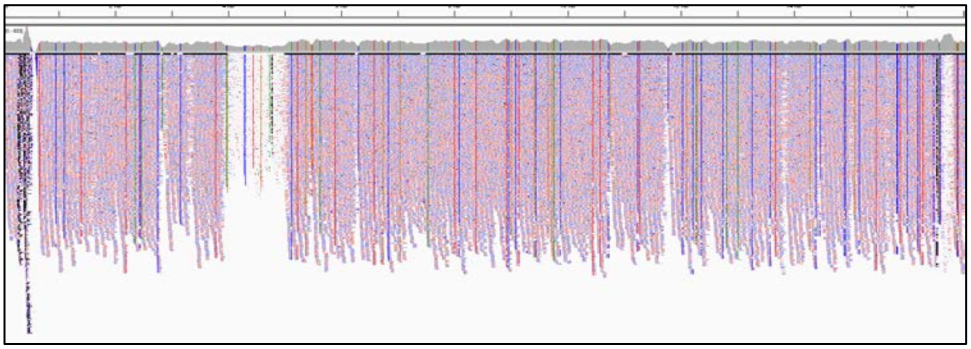

Historical Lion AMNH_81840 with numt correction

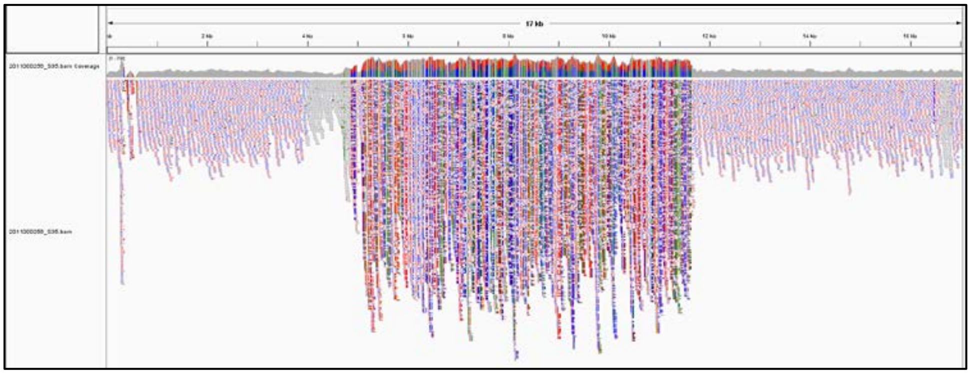

Modern Lion 2011000256 without numt correction

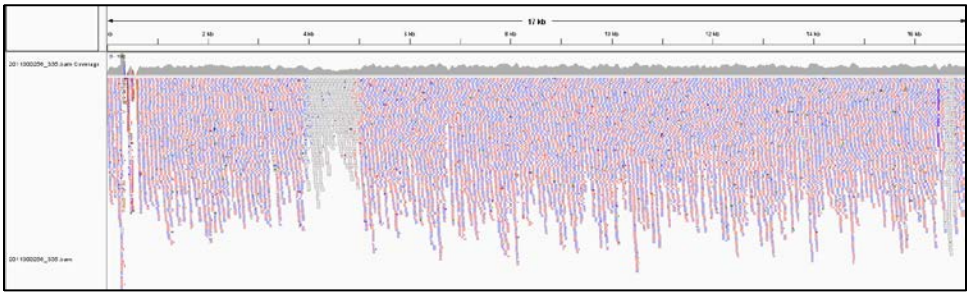

Modern Lion 2011000256 with numt correction

Whole genome sequence coverage ranged from 1.3x-9.6x with mitochondrial coverage ranging from 1.9x-1703.9x (See Supporting Information S1.d).

***Mitogenome Filtering and SNP Identification***

The mitogenome was isolated from genomic reads and filtered to produce a panel of polymorphic sites for further analysis. Conservative filtering was implemented to accommodate the higher error rate associated with DNA damage possible in older samples (9–11). BAM files were filtered with a mismatch threshold of 0.03 using SAMtools (12). Variants were called for the Mitogenome via GATK (13) HaplotypeCaller tool then filtered for a minimum depth of 10-bp, retention of only biallelic SNPs, and removal of heterozygotes using VCFtools (14) and BCFtools (12). FASTA sequences were created for individuals containing <10% missing data across the mitogenome using BCFtools. The mitogenome of modern sample M_ZAM_T256 was annotated using the MITOS webserver (17). The annotation of M_ZAM_T256 mitogenome can be found in Supporting Information S12.

The resulting sequences were combined with published modern mitogenomes from GenBank (KP001493- KP001506 (15), KP202262 (8), KC834784 (16)) and aligned using the EMBL-EBI web tool Clustal Omega (www.ebi.ac.uk/Tools/msa/clustalo). Polymorphic sites were identified using dnaSP (18). To account for potential biases produced by differences in sequencing methods between studies, the multiple sequence alignment (MSA) was trimmed to include only polymorphic sites present in mitogenome sequences generated in this study.

### S11. Annotation of Lion Mitogenome (Sample: M-ZAM-T256)

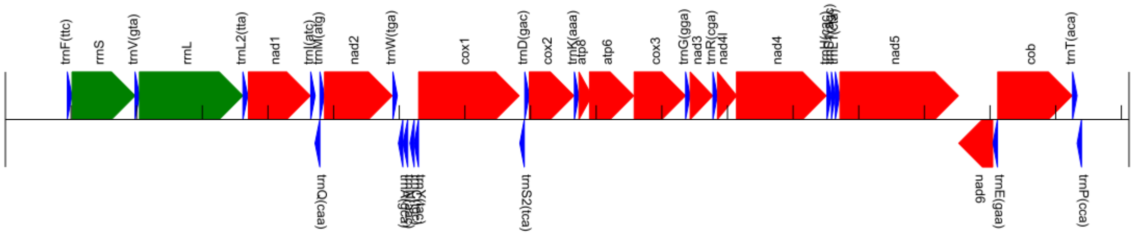


| Abbreviation | Name | Seq | Direction | Start | End |
|---|---|---|---|---|---|
| tRNA-Phe | Phenylalanine | trnF(ttc) | + | 943 | 1015 |
| 12S | ribosomal RNA | rrnS | + | 1015 | 1977 |
| tRNA-Val | Valine | trnV(gta) | + | 1975 | 2044 |
| 16S | ribosomal RNA | rrnL | + | 2042 | 3618 |
| tRNA-Leu2 | Leucine 2 | trnL2(tta) | + | 3618 | 3693 |
| ND1 | NADH dehydrogenase subunit 1 | nad1 | + | 3701 | 4646 |
| tRNA-Ile | Isoleucine | trnI(atc) | + | 4651 | 4720 |
| tRNA-Gln | Glutamine | trnQ(caa) | - | 4717 | 4791 |
| tRNA-Met | Methionine | trnM(atg) | + | 4792 | 4861 |
| ND2 | NADH dehydrogenase subunit 2 | nad2 | + | 4861 | 5893 |
| tRNA-Trp | Tryptophan | trnW(tga) | + | 5903 | 5972 |
| tRNA-Ala | Alanine | trnA(gca) | - | 5987 | 6056 |
| tRNA-Asn | Asparagine | trnN(aac) | - | 6057 | 6130 |
| tRNA-Cys | Cysteine | trnC(tgc) | - | 6163 | 6228 |
| tRNA-Tyr | Tyrosine | trnY(tac) | - | 6228 | 6294 |
| COI | cytochrome c oxidase subunit I | cox1 | + | 6295 | 7828 |
| tRNA-Ser2 | Serine 2 | trnS2(tca) | - | 7837 | 7906 |
| tRNA-Asp | Aspartic acid | trnD(gac) | + | 7912 | 7981 |
| COX2 | cytochrome c oxidase subunit II | cox2 | + | 7981 | 8662 |
| tRNA-Lys | Lysine | trnK(aaa) | + | 8668 | 8736 |
| ATP8 | ATP synthase F0 subunit 8 | atp8 | + | 8737 | 8935 |
| ATP6 | ATP synthase F0 subunit 6 | atp6 | + | 8898 | 9573 |
| COX3 | cytochrome c oxidase subunit III | cox3 | + | 9578 | 10361 |
| tRNA-Gly | Glycine | trnG(gga) | + | 10362 | 10431 |
| ND3 | NADH dehydrogenase subunit 3 | nad3 | + | 10431 | 10776 |
| tRNA-Arg | Arginine | trnR(cga) | + | 10778 | 10847 |
| ND4l | NADH dehydrogenase subunit 4L | nad4l | + | 10847 | 11141 |
| ND4 | NADH dehydrogenase subunit 4 | nad4 | + | 11137 | 12505 |
| tRNA-His | Histidine | trnH(cac) | + | 12515 | 12584 |
| tRNA-Ser1 | Serine 1 | trnS1(agc) | + | 12584 | 12643 |
| tRNA-Leu1 | Leucine 1 | trnL1(cta) | + | 12643 | 12713 |
| ND5 | NADH dehydrogenase subunit 5 | nad5 | + | 12713 | 14519 |
| ND6 | NADH dehydrogenase subunit 6 | nad6 | - | 14523 | 15042 |
| tRNA-Glu | Glutamic acid | trnE(gaa) | - | 15045 | 15114 |
| cytb | cytochrome b | cob | + | 15117 | 16251 |
| tRNA-Thr | Threonine | trnT(aca) | + | 16257 | 16327 |
| tRNA-Pro | Proline | trnP(cca) | - | 16327 | 16394 |
| D-loops | control region | | | | |

>ZAM_T256; 944-1015; +; trnF(ttc)
GTTAATGTAGCTTAAACACATTTAAAGCAAGGCACTGAAAATGCCTAGATGAGTCGCCAGACTCCATAAACA

>ZAM_T256; 1016-1977; +; rrnS
CAAAGGTTTGGTCCTAGCCTTTCCATTAGTTATTAATAAAATTACACATGCAAGCCTCCGCATCCCGGTGAAAATGCCCTCTAAA
TCACCTAGTGATCCAAAGGAGCTGGTATCAAGCACACAACCATTGTAGCTCACAACACCTTGCTCAGCCACACCCCCACGGGAT
ACAGCAGTGATAAAAATTAAGCTATGAATGAAAGTTCGACTAAGCTATATTAAACTAGGGTTGGTAAATTTCGTGCCAGCCAC
CGCGGTCATACGATTAACCCAGACTAATAGACTTACGGCGTAAAGCGTGTTACAGAAGAAAAATATACTAAAGTTAAACCTTA
ACTAGGCTGTAAAAAGCTGCAGTTAACATAAAAATACAGCACGAAAGTAACTTTAATACCTCCGACCACACGATAGCTAAGAT
CCAAACTGGGATTAGATACCCCACTATGCTTAGCCCTAAACCTAGATAGTTAACCCAAACAAAACTATCCGCCAGAGAACTACT
AGCAACAGCTTAAAACTCAAAGGACTTGGCGGTGCTTTACATCCCTCTAGAGGAGCCTGTTCTATAATCGATAAACCCCGATAA
ACCTCACCATCTCTTGCTAATTCAGCCTATATACCGCCATCTTCAGCAAACCCTAAAAAGGAAGAAAAGTAAGCACAAGTGTCT
TAACACAAAAAAGTTAGGTCAAGGTGTAGCCTATGAGATGGGAAGCAATGGGCTACATTTTCTACAATTAGAACACCCACGAA
AATCCTTATGAAACTAAGCATTCAAGGAGGATTTAGCAGTAAATTTGAGAATAGAGAGCTCAATTGAATCGGGCCATGAAGCA
CGCACACACCGCCCGTCACCCTCCTCAAGTGACTAGCCCCTAAAGAAACCTATTCAAACCACTACATCCACAAGAGGAGACAAG
TCGTAACAAGGTAAGCATACTGGAAAGTGTGCTTGGATGACA

>ZAM_T256; 1976-2044; +; trnV(gta)
CAAGATGTAGCTTAAACTAAAGCGTCTGGCTTACACCCAGAAGATTTCATATTAAACTGACCGTCTTGA

>ZAM_T256; 2043-3618; +; rrnL
GAGCCAAAGCTAGCCCAATCATCTACAAACGCAACTAACACTAGAAAGTAAAATAAAACATTTAGTTACCCCATAAAAGTATA
GGAGATAGAAATTTAACTTGGCGCTATAGAGAAAGTACCGCAAGGGAAGGATGAAAGAAAAAACTAAAAGCACTATACAGC
AAAGATTGCCCCTTGTACCTTTTGCATAATGAGTTAGCTAGTAACAGCCTAACAAAGAGAACTTCAGCTAGGCCCCCCGAAACC
AGACGAGCTACCCATGAACAATCTATTACAGGATGAACTCGTCTATGTTGCAAAATAGTGAGAAGATTTGTGGGTAGAGGTGA
AAAGCCTAACGAGCCTGGTGATAGCTGGTTGCCCAGAACAGAATCTTAGTTCAACTTTAAACTTACCTCAAAACCCTAAAATTC
CAATGTAAGTTTAAATTATAGTCTAAAAAGGTACAGCTTTTTAGAACTAGGATACAGCCTTAATTAGAGAGTAAGCACAAACAC
AAACCATAGTTGGCCTAAAAGCAGCCACCAATTAAGAAAGCGTTCAAGCTCGACAATCAAAACATCTCAATGTCAAAAAACGT
AACCAACTCCTAACCTAAAACTGGGCTAATCTATTTAATAATAGAAGCAATAATGCTAATATGAGTAACAAGAAGCATTTCTCC
CGTGCATAAGCTTATATCAGAACGGATAACCACTGATAGTTAACAACAAGATAGATACAACCTAACTACAAGCAAAATATCAA
ACTAATTGTTAACCCAACACAGGCATGCAATCCAGGGAAAGATTAAAAGAAGTGAAAGGAACTCGGCAAACACAAGCCCCGC
CTGTTTACCAAAAACATCACCTCTAGCATTTCCAGTATTAGAGGCACTGCCTGCCCAGTGACATTAGTTAAACGGCCGCGGTAT
CCTGACCGTGCAAAGGTAGCATAATCATTTGTTCCTTAAATAGGGACTTGTATGAATGGCCACACGAGGGCTTTACTGTCTCTT
ACTTCTGATCCGTGAAATTGACCTTCCCGTGAAGAGGCGGGAATATGACAATAAGACGAGAAGACCCTATGGAGCTTTAATTA
ACCGACCCAAAGAGATCTTGATAATCAACCAACAGGGATAACAAACCTCTACCATGGGTCGACAATTTAGGTTGGGGTGACCT
CGGAGAATAAAACAACCTCCGAGTGATTTAAATCTAGACTAACCAGTCGAAAATATTACATCACTTATTGATCCAAAAACTTGA
TCAACGGAACAAGTTACCCTAGGGATAACAGCGCAATCCTATTTTAGAGTCCATATCGACAATAGGGTTTACGACCTCGATGTT
GGATCAGGACATCCCGATGGTGCAGCAGCTATCAAAGGTTCGTTTGTTCAACGATTAAAGTCCTACGTGATCTGAGTTCAGAC
CGGAGTAATCCAGGTCGGTTTCTATCTATTAAATAATTTCTCCCAGTACGAAAGGACAAGAGAAATAAGGCCCACTTTACCAAA
GCGCCTTTAACCAAATAGATGATATAATCTCAATCTAGACAGTTTATCTAAACACATCGCCCGAGAGCTCGGGTTT

>ZAM_T256; 3619-3693; +; trnL2(tta)
GTTAGGGTGGCAGAGCCCGGTAATTGCATAAAACTTAAGCTTTTATCATCAGAGGTTCAACTCCTCTCCCTAACA

>ZAM_T256; 3702-4646; +; nad1
ATAATCAATATCCTCTCACTAATCATCCCCATTCTCCTCGCCGTAGCCTTCCTAACCCTAGTTGAACGTAAAGTACTAGGCTACAT
ACAACTTCGCAAAGGACCAAATGTCGTAGGGCCATATGGCCTACTTCAACCCATTGCAGACGCCATAAAACTCTTCACTAAAGA
ACCCCTCCGGCCCCTTACATCCTCTACATTCATATTTATTATAGCACCTATCCTAGCCCTTACACTAGCCCTAACCATATGAATCC
CACTGCCCATACCATATCCACTCGTTAACATAAACCTAGGGGTGCTATTCATACTAGCCATATCCAGCCTAGCTGTTTACTCCAT
CCTATGATCCGGGTGGGCTTCAAACTCAAAATATGCTCTAATCGGTGCCCTACGAGCCGTAGCTCAAACAATCTCATACGAAGT
CACACTAGCTATTATCCTCTTATCAGTACTACTAATAAACGGATCCTTCACATTAGCCACACTAATCACCACCCAAGAATATATCT
GACTTATTATCCCCGCATGACCCCTAGCTATAATATGATTCATCTCTACACTAGCAGAGACCAACCGAGCCCCATTTGACCTCAC
AGAAGGAGAATCAGAGCTTGTCTCCGGATTTAACGTAGAATACGCAGCAGGTCCTTTCGCCCTATTCTTTTTAGCAGAATATGC
CAACATTATCATAATAAACATCCTCACAACAATCCTGTTCTTCGGAGCATTTCATAGTCCCTACATACCAGAACTATATACCACTA
ACTTCACCGTAAAAACCCTGATCTTAACAACCACCTTCCTATGAATCCGAGCATCTTATCCACGATTCCGATACGACCAACTAAT
ACACCTCCTATGAAAAAGCTTTCTACCCCTTACCCTAGCTCTATGTATATGGCACGTCTCCCTACCCATTATCACAGCAAGTATCC
CACCTCAA

>ZAM_T256; 4652-4720; +; trnI(atc)
AGAAATATGTCTGATAAAAGAATTACTTTGATAGAGTAAAACATAGAGGTTTAAGCCCTCTTATTTCTA

>ZAM_T256; 4718-4791; -; trnQ(caa)
TAGAATGTGGTGTAATATTGGTAGCACGAAGATTTTTGGATTCTTAGGATTAGGTTCGACTCCTATAATTCTAG

>ZAM_T256; 4793-4861; +; trnM(atg)
AGTAAGGTCAGCTAAATAAGCTATCGGGCCCATACCCCGAAAATGTTGGTTTATACCCTTCCCATACTA

>ZAM_T256; 4862-5893; +; nad2
ATCAAACCCCCCATTTTTATTATCATTATATTAACCGTTATCTCAGGAACCATAATCGTAATAACAGCCTCCCACTGACTTATAGT
CTGAATCGGCTTTGAAATAAACCTACTAGCCATCATTCCCATCCTCATAAAAAAATACAACCCACGAGCCACAGAAGCAGCCAC
AAAATATTTCCTAACACAAGCAACCGCTTCAATACTCCTAATAATAGGAATCATTATCAACTTACTGCACTCAGAACAATGAACC
GTATCAAAGGATCTTAACCCCATAGCATCCATCGTAATAACAACCGCCCTAGCAATAAAACTAGGACTAGCCCCATTCCACTTCT
GAGTACCCGAAGTTACACAAGGAATCCCCATATCCTCGGGCCTAATCCTACTCACATGACAAAAAATCGCCCCACTATCAATCC
TATACCAAATCTCACCCACCATCAACCCCAACCTACTCCTAACAATAGCTACCATATCAGTTATAATTGGAGGCTGAGGAGGAC
TTAATCAGACCCAACTACGAAAAATCATAGCATATTCCTCAATTGCCCACATAGGCTGAATAGCAGCCATCATAATATACAGCC
CCACAATAATAATTTTAAACCTAACCATCTACATCACCATAACACTAACCACCTTCATACTATTCATATACAACTCCACCACAACA
ACATCATCCCTATCGCAAACATGAAACAAAACACCCCTAATCACCTCATTTATTCTAGTGCTAATAATATCTCTAGGCGGCCTTC
CTCCGCTCTCCGGCTTTATCCCAAAATGAATAATCATTCAAGAACTAACTAAAAATGAAATAATTATAATACCCACACTACTAGC
TATAACAGCACTACTTAACCTGTACTTCTACATACGACTAACATACACCACTGCACTAACTATATTCCCCTCAAACAACTGCATAA
AAATAAAATGACGATTCGAGCGCACAAAAAAAACAATCCTTTTGCCCCCCTTAATTGTAATATCTACTATACTACTACCACTCAC
ACCGATACTATCT

>ZAM_T256; 5904-5972; +; trnW(tga)
AGAAGTTTAGGTTAAACTAGACCAAGAGCCTTCAAAGCTCTAAGCAAGCCCTAACAGACTTAACTTCTG

>ZAM_T256; 5988-6056; -; trnA(gca)
GAGGGCTTAGCTTAATTAAAGTGTTTGATTTGCAATCAATTGATGTAAGATAGATTCTTGCAGTCCTTA

>ZAM_T256; 6058-6130; -; trnN(aac)
TAGATTGAAGCCAGTTGATTAGGGTATTTAGCTGTTAACTAAAATTTCGTGGGGTTGGGGCCCACCAATCTAG

>ZAM_T256; 6164-6228; -; trnC(tgc)
AGTCCTGCAGTGAATATCATATTGAATTGCAAATTCAAAGAAGCAGCTTGACGCTGCCGGGGCTT

>ZAM_T256; 6229-6294; -; trnY(tac)
GGTAAAATGGCTGAGTAAGCATTAGACTGTAAATCTAAAGACAGAGGTTCAGTCCTCTTTTTACCA

>ZAM_T256; 6296-7828; +; cox1
ATGTTCATAAATCGCTGACTATTTTCAACCAATCACAAAGATATTGGAACTCTTTACCTTCTATTTGGTGCCTGGGCTGGTATGG
TGGGGACTGCTCTCAGTCTCCTAATCCGAGCCGAACTGGGTCAACCTGGCACACTACTAGGGGATGACCAAATTTATAATGTA
GTCGTCACCGCCCATGCTTTTGTAATAATCTTCTTTATAGTAATACCTATCATGATTGGAGGATTCGGAAACTGATTGGTCCCAT
TAATAATTGGAGCCCCCGATATAGCATTCCCTCGAATGAACAATATAAGCTTCTGACTTCTTCCCCCGTCTTTCCTACTTTTGCTT
GCATCATCTATGGTAGAAGCTGGAGCAGGAACTGGGTGGACAGTATACCCGCCTCTAGCCGGCAACCTAGCTCACGCAGGAG
CATCTGTAGATCTAACTATTTTTTCACTACACCTGGCAGGTGTCTCCTCAATCCTAGGTGCTATTAATTTTATTACTACTATTATTA
ATATAAAACCCCCTGCTATATCCCAATACCAAACACCTTTATTTGTCTGATCGGTTTTAATCACTGCTGTATTGCTACTCCTATCA
CTGCCAGTTTTAGCAGCAGGCATCACTATGCTACTGACAGATCGAAATCTGAATACCACATTTTTTGACCCTGCCGGAGGAGGG
GACCCTATCTTATACCAACATCTATTCTGATTTTTTGGTCACCCAGAAGTCTACATTTTAATTTTACCCGGATTCGGAATAATTTC
ACACATTGTCACCTATTATTCAGGTAAAAAAGAACCTTTTGGCTACATAGGAATAGTTTGAGCTATAATATCAATTGGTTTTCTG
GGCTTTATTGTGTGAGCCCATCACATGTTTACTGTGGGGATAGATGTGGACACACGAGCATACTTTACATCTGCTACTATAATT
ATTGCTATTCCCACTGGAGTAAAAGTATTTAGCTGACTGGCCACTCTTCATGGCGGTAATGTCAAATGGTCTCCCGCTATGCTG
TGAGCCCTAGGATTCATCTTCCTATTTACTGTTGGGGGCTTAACAGGAATTGTACTAGCAAATTCCTCATTAGATATTGTCCTTC
ACGATACATACTATGTAGTAGCCCACTTCCACTATGTATTGTCGATAGGAGCAGTATTTGCTATTATAGGGGGCTTCGTTCATT
GATTCCCCCTATTCTCAGGGTATACTCTCGATAATACCTGGGCAAAAATTCATTTTACGATTATGTTCGTAGGCGTCAATATAAC
GTTTTTCCCTCAGCATTTTCTAGGCTTGTCCGGAATGCCTCGACGTTATTCTGACTACCCAGACGCATATACAACTTGAAACACA
GTCTCCTCAATAGGCTCTTTTATTTCATTAACAGCAGTAATATTAATGGTTTTCATAGTGTGAGAGGCTTTTGCATCAAAGCGAG
AAGTGGCCATAGTGGAACTAACCACGACTAATCTTGAATGACTACATGGATGTCCCCCTCCATACCACACATTTGAAGAACCAA
CCTATGTG

>ZAM_T256; 7838-7906; -; trnS2(tca)
GAGAAAGACATAGTGGTTATGAAATTGGCTTGAAACCAGTCTGAGGAGGTTCGATCCCTTCCTTTCTTA

>ZAM_T256; 7913-7981; +; trnD(gac)
GAAGTATTAGTAAAACAATTACATAACTTTGTCAAGGTTAAATTATAGGTTTAAGCCCTGTGTACTTCC

>ZAM_T256; 7982-8662; +; cox2
ATGGCATACCCCTTCCAGCTAGGTCTCCAAGATGCTACATCCCCCATTATAGAAGAGCTTCTACACTTCCACGATCACACGTTAA
TAATTGTATTTCTAATTAGTTCCCTAGTCCTTTATATCATCTCACTAATGCTGACAACTAAACTCACGCACACAAGTACAATAGAT
GCCCAAGAAGTAGAAACCATCTGAACTATTTTACCAGCAATTATCTTAATTCTCATTGCCCTGCCCTCCTTACGAATTCTCTATAT
AATAGACGAGATTAATAGTCCCTCTCTCACTGTAAAGACCATGGGACATCAGTGATATTGAAGCTATGAGTATACTGACTATGA
AGACCTGAGCTTTGACTCCTATATAATCCCTACTCAAGAGCTAAAGCCCGGAGAACTCCGACTATTAGAAGTCGATAACCGAGT
AGTATTGCCAATAGAAGTAACTGTTCGCATGTTAATCTCATCAGAAGACGTGTTGCACTCATGAGCCGTTCCATCCCTAGGTCT
AAAAACTGACGCTATTCCAGGCCGACTAAACCAAACAACCCTAATGGGTACACGACCTGGACTATACTATGGTCAATGCTCAG
AGATCTGCGGCTCAAACCACAGTTTTATGCCTATTGTCCTTGAACTAGTCCCACTGTCATACTTTGAAAAGTGATCTGTGTCTAT
ACTG

>ZAM_T256; 8669-8736; +; trnK(aaa)
CATTAAGAAGCTAAATTAGCGTTAACCTTTTAAGTTAAAAACTGGGAGTTTAGACCTCCCCTTAATGG

>ZAM_T256; 8738-8935; +; atp8
ATGCCACAGTTAGATACATCAACCTGATTCATTACCATTATTTCAATAATTATAACACTATTTATTATATTTCAATTAAAAATCTC
AAAACACTTATACCCATCGAGCCCGGAGCCTAAATCTACAGCTGCACTAAAACAACCTAATCCTTGAGAAAAAAAATGAACGA
AAATCTATTCACCTCTTTCACTACCCCAA

>ZAM_T256; 8899-9573; +; atp6
ATGAACGAAAATCTATTCACCTCTTTCACTACCCCAACAATAATAGGACTGCCTGTTGTCGTATTAATTATTATGTTCCCCAGCAT
TCTATTCCCCTCACCCAACCGACTAATTAATAACCGCCTAGTCTCACTCCAACAATGATTAGTACAACTAACATCAAAACAAATA
TTAGCTATTCACAATCACAAAGGACAAACCTGAGCCCTAATACTCATGTCTCTCATTTTATTTATCGGATCCACAAACCTGTTGG
GCCTACTGCCCCACTCATTTACCCCAACTACCCAATTATCAATAAACTTAGGAATAGCTATCCCTCTATGAGCCGGTACCGTAGT
CACCGGATTTCGCCACAAAACTAAAGCGTCCCTGGCTCACTTTCTACCACAAGGAACACCAATCCCCTTAATTCCTATGCTTGTA
ATTATTGAAACCATTAGCCTTTTTATTCAACCCGTGGCTCTGGCCGTACGACTTACAGCCAACATTACTGCGGGTCACTTATTAA
TACACTTAATTGGAGGAGCTACCCTGGCTCTGACAAACATTAATGCCTCTGTCGCTTTAATTACCTTTATCATCCTCATCCTGCTG
ACAGTCCTTGAATTCGCTGTGGCCCTAATCCAAGCCTACGTCTTTACCCTACTTGTAAGCCTGTACCTACATGACAAT

>ZAM_T256; 9579-10361; +; cox3
ATGACCCATCAAACCCACGCATACCATATGGTTAACCCCAGCCCATGACCACTTACGGGGGCTCTCTCAGCCCTACTGATAACC
TCAGGTCTGGCTATATGATTTCACTACAACTCAACATTATTATTAACCCTAGGTATAACCACCAACCTACTGACTATATATCAAT
GGTGACGAGATATTATTCGGGAAAGCACATTCCAAGGTCACCACACGCCTATCGTTCAAAAAGGTCTCCGTTATGGAATAGTT
CTCTTTATCATCTCGGAAGTATTCTTCTTCGCAGGCTTTTTCTGGGCCTTCTATCACTCAAGCCTGGCCCCAACCCCCGAATTAGG
AGGATGCTGGCCACCAACAGGTATTATTCCCCTAAACCCCCTAGAAGTCCCACTGCTTAACACTTCCGTACTTTTAGCCTCCGGA
GTATCAATTACCTGAGCCCACCATAGTTTAATGGAAGGCAATCGAAAACATATACTCCAAGCACTATTTATTACAATCTCCCTAG
GGGTCTACTTTACCCTCCTCCAAGCCTCCGAATACTATGAAACATCATTTACAATCTCAGACGGGGTCTATGGATCCACCTTCTT
CATAGCTACAGGATTCCACGGCCTACACGTAATTATTGGCTCTACCTTCTTAATTGTATGTTTCTTGCGCCAACTAAAATATCATT
TCACATCGAGCCACCATTTTGGATTTGAAGCCGCTGCTTGATATTGACACTTCGTAGATGTGGTTTGACTATTCCTATACGTTTC
CATTTATTGATGAGGATCC

>ZAM_T256; 10363-10431; +; trnG(gga)
ATTCCTTTAGTATCAACAAGTACAGCTGACTTCCAATCAGCCAGTTTCGGTAGAATCCGAAAAGGAATA

>ZAM_T256; 10432-10776; +; nad3
ATAAACATAATAATTGCCCTGCTCACCAATACACTTCTATCCACACTACTTGTATTTATTGCATTCTGACTGCCCCAACTAAACAT
TTACGCAGAAAAAGCAAGTCCCTATGAATGTGGATTTGACCCCATAGGATCCGCCCGCCTACCCTTCTCCATAAAATTTTTCTTA
GTAGCTATTACATTCTTACTATTTGATCTAGAAATTGCACTACTACTCCCTCTTCCTTGGGCCTCACAAACAAACAAATTACCAAC
CATACTCATCACAGCCCTCCTACTAATCTCTCTACTAGCCGTAAGCCTAGCCTACGAATGAACCCAAAAAGGACTAGAATGAAC
TGAA

>ZAM_T256; 10779-10847; +; trnR(cga)
TGATAATTAGTTTAAACTAAAACAAATGATTTCGACTCATTAGATTGTAGCTTACCCTATAATTATCAG

>ZAM_T256; 10848-11141; +; nad4l
ATGTCCATGGTCTATGTTAATATATTTCTGGCTTTCATCATGTCACTCATAGGACTATTAATGTACCGATCCCATTTAATATCATC
CCTCCTATGTCTAGAAGGCATAATACTATCCCTATTCATCATAATAACCGTGACAATTCTAAATAATCATTTCACACTAGCTAGC
ATGACTCCCATCATTCTGCTAGTATTCGCAGCCTGCGAGGCGGCACTGGGCTTATCCTTACTAGTAATGGTATCAAACACATAC
GGTACCGACTATGTACAAAACCTAAACCTCCTACAATGC

>ZAM_T256; 11138-12505; +; nad4
ATGCTAAAAATTATTATCCCCACTGCCATACTCATACCAATAACATGACTATCAAAACCCAACATAATTTGAATTAACTCAACTA
CCTACAGCCTTCTGATCAGCCTTATTAGTCTCCCCTACTTAAACCAACTAGGCGACAACAGCCTGAATCTCTCATTACTATTTTTC
TCAGACTCACTTTCTGCACCTCTACTAGTCTTAACAACATGGCTCCTACCACTAATGCTCATGGCTAGTCAATCCCACCTGTCAAA
AGAGACCTTGGCCCGAAAAAAACTATACATCACAATACTTATTATTTTGCAACTTCTCCTAATTATAACATTCACTGCCACAGAA
TTAATCATATTCTACATTCTATTTGAAGCCACATTAATCCCCACTCTCATCATCATTACTCGATGGGGTAATCAAACAGAGCGAC
TAAACGCTGGTCTATACTTTCTATTCTACACCCTGATAGGCTCACTGCCCCTCCTAGTCGCACTATTATATATTCAAAACACAACA
GGGACTTTAAATTTCCTAATCATCCAATACTGAGCCAAACCAATCTCAGCCACCTGATCTAACATCTTCCTCTGACTAGCATGCA
TAATAGCATTTATAGTAAAAATACCTTTATACGGACTCCACCTGTGGTTACCAAAAGCACACGTCGAAGCCCCCATTGCCGGCT
CAATAGTACTTGCCGCTGTACTATTAAAACTGGGGGGATATGGAATGATACGCATTACAATTTTACTAAACCCCATAACAAACC
AAATAGCATACCCCTTCATAATGCTATCCCTATGAGGAATAGTCATGACAAGTTCTATCTGTCTACGCCAGACGGACCTGAAAT
CCTTAATCGCATACTCATCAGTAAGCCACATAGCCCTAGTAATTGTAGCTGTACTAATCCAAACGCCCTGAAGCTATATAGGAG
CTACAGCCCTTATAATCGCTCACGGACTAACCTCCTCAATACTATTCTGTCTTGCAAACTCAAACTACGAACGAGTCCATAGCCG
AACAATAATTCTAGCACGAGGCCTGCAAACCATCCTCCCTCTAATAGCTGCCTGATGATTACTAGTCAGCCTCGCGAACCTAGC
CCTGCCCCCTACCATTAACCTAATTGGAGAACTATTCGTAGTGATAGCCTCCTTCTCATGATCTAACATAACTATCGTCCTCATAG
GCACAAACATTATCATCACAGCCTTATATACCCTCTACATACTCACTACAACCCAACGAGGCAAGTATACACACCATATCAAAAA
TATCAATCCATCATTCACACGAGAAAACGCCCTAATAGCCCTTCACCTACTCCCACTCCTTCTCTTATCCCTCAACCCCAAAATCG
TACTAGGC

>ZAM_T256; 12516-12584; +; trnH(cac)
GTAAATATAGTTTAACAAAAACATTAGATTGTGAATCTAACAATGGAAGTGCAAATCTTCTTATTTACC

>ZAM_T256; 12585-12643; +; trnS1(agc)
GAAAAAGTATGCAAGAACTGCTAATTCATGCCCCCACGTATAAAAACGTGGCTTTTTCA

>ZAM_T256; 12644-12713; +; trnL1(cta)
ACTTTTATAGGATAGAAGTAATCCATTGGCCTTAGGAGCCAAAAAATTGGTGCAACTCCAAATAAAAGTA

>ZAM_T256; 12714-14519; +; nad5
ATAAACCTATTTGCCTCCTTTATACTCACTTCAATATTTATGCTACTCCTACCTATTATTATATCCAACACTCAACTATACAAAAAC
AACCTATACCCCCACTATGTGAAAACCACAATCTCTTATGCCTTCGCCATCAGCATAATTCCGGCCATAATATTCGTCTCCTCCG
GACAAGAAACAATCGTCTCAAACTGACACTGACTGTCAATTCAAACTCTCAAGTTGTCACTAAGCTTTAAACTAGATTATTTCTC
GATCATTTTCATCCCTGTAGCACTTTTCGTTACATGGTCGATCATGGAGTTCTCAATATGGTACATGCACACAGATCCTTATATTA
ACCGATTCTTCAAGTATCTCCTCATATTTCTAATCACCATAATAATCCTAGTAACCGCCAATAACCTGTTCCAACTGTTTATTGGT
TGAGAAGGAGTAGGAATCATATCCTTTCTACTCATTGGATGATGATATGGTCGAGCAGACGCAAACACTGCCGCCCTACAAGC
AATTCTCTATAACCGCATCGGAGATGTAGGATTTATCACGGCTATAGCATGATTCCTCGCCAACATAAATGCATGAGACTTCCA
ACAAATCTTTATTACCCAACACAAAAACCTAAATATTCCACTACTAGGACTTCTTCTAGCAGCCACAGGCAAGTCTGCCCAATTT
GGCCTACATCCATGACTACCATCAGCCATAGAGGGTCCAACCCCTGTCTCTGCCCTACTCCACTCAAGCACAATAGTTGTAGCC
GGAGTCTTTTTATTAATCCGCTTCCACCCACTCATAGAACAAAACAAAACCATACAGACCCTCACTCTATGCCTAGGAGCCATCA
CAACCCTATTCACAGCCATCTGTGCTCTCACACAAAATGATATCAAAAAAATCGTTGCCTTCTCAACCTCAAGCCAATTGGGTCT
AATAATCGTCACTATCGGAATCAACCAACCCTACCTCGCATTCCTCCATATCTGTACACATGCATTTTTCAAAGCCATACTATTCA
TATGCTCTGGGTCAATTATTCATAGCCTAAATGATGAACAAGACATTCGAAAAATAGGCGGACTATACAAACCAATACCCTTCA
CCACCTCCTCCCTCATCATCGGAAGTCTCGCATTAACAGGTATACCTTTCCTAACAGGCTTTTACTCCAAAGACCTAATCATCGA
GACAGCCAATACGTCGTATACCAACGCCTGAGCCCTGTCGGTCACTCTCATTGCTACATCTCTTACGGCCGCCTATAGTACTCGA
ATCATATTCTTTGCACTTCTAGGACAGCCCCGGTTTAACTCCCTAAGTCCAATCAATGAAAACAACCCCCACCTTATCAACTCCAT
TAAACGTCTCTTAGTTGGAAGCATTTTTGCAGGATACTTGATTTCCCATAATATCCCCCCAACAACCATCCCACAAATAACTATG
CCCCACTATCTAAAGCTTACTGCCCTTGCCATAACCATTACAGGCTTCATCTTAGCATTAGAACTTAATCTCGCAGCTAAAAACTT
AAAATTTAAATATCCCTCAAACCTCTTTAAGTTTTCTAACCTCCTAGGGTACTTTCCAACTGTAATACACCGCCTCCCATCAACAA
TAAGCCTAACTATAAGCCAAAAATCTGCATCGATACTATTAGATATAATCTGGCTAGAAAGTGTATTACCAAAATCTATCTCCCA
CTTCCAAATAAAAATATCAACCATTGTATCTAATCAGAAAGGACTAGTTAAACTTTACTTCTTATCCTTCATAATCACCTTGACCC
TTAGCCTACTCCTACTT

>ZAM_T256; 14524-15042; -; nad6
ATAACATATATTGTATTTATTTTAAGTACAATTTTCGTGGTGAGCTTTGTGAGTTTTTCTTCAAAGCCCTCTCCTATTTATGGTGG
GTTTGGTTTGATTGTGGCTGGTGGTGTTGGTTGCGGTATTGTACTGAATTTTGGAGGATCGTTTTTAGGTTTAATGGTCTTTTTA
ATTTATTTGGGGGGTATACTTGTGGTGTTTGGGTACACTACAGCTATGGCTACTGAGCCTTATCCTGAGGCATGGACGTCTAAT
AAGGCTGTGTTGGGTGCATTTATTACGGGAGTGCTAGCGGAATTATTAACTGCTTGCTATATTTTAAAAGAAGACGAGGTTGA
AGTTGTGTTTAAGTTTAATGGTGCAGGTGATTGGGTGATTTATGATACGGGTGATTCAGGATTTTTTAGTGAGGAGGCCATGG
GAATTGCAGCACTGTATAGCTATGGAACCTGGTTGGTAGTTGTTACTGGTTGGTCCTTGCTTATTGGTGTATTGGTGATTATAG
AGGTTACTCGTGGA

>ZAM_T256; 15046-15114; -; trnE(gaa)
GTTCTTATAGTTGAAATACAACGGTGGTTTTTCATATCATTAGTCATGGTTAGATTCCATGTGAGAATT

>ZAM_T256; 15118-16251; +; cob
ATGACCAACATTCGAAAATCACACCCCCTTGTCAAAATTATTAATCACTCATTCATTGATCTTCCCACTCCACCCAATATCTCAGC
ATGATGAAACTTTGGCTCCTTATTAGGAGTATGTTTAATCCTACAAATTCTCACCGGCCTCTTTCTAGCCATACATTACACACCA
GACACAATAACCGCTTTCTCATCAGTCACCCACATTTGCCGCGATGTAAACTATGGCTGAATTATCCGGTACCTACACGCCAAC
GGAGCCTCCATATTCTTTATCTGCCTATACATGCATGTAGGACGAGGAATATACTATGGCTCCTATACTTTCTCAGAAACATGAA
ACATTGGAATCATATTGTTGCTCACAGTTATAGCTACAGCCTTCATAGGATATGTCTTACCGTGGGGCCAAATATCCTTTTGAGG
TGCAACTGTAATCACTAATCTCCTATCAGCAATCCCATACATCGGGGCCGACCTAGTAGAGTGGATCTGAGGAGGCTTCTCAGT
AGACAAAGCCACCCTGACACGATTCTTTGCCTTCCACTTCATCCTTCCATTTATCATCTCAGCCCTAGCAGCAGTCCACCTCCTAT
TCCTCCATGAAACAGGATCTAATAACCCCTCAGGAATGGTATCTGACTCAGATAAAATTCCATTCCATCCATACTATACAATCAA
AGATATCCTAGGCCTTCTAGTACTAATCTTAACACTCATACTACTCGTCCTATTCTCACCAGACCTATTAGGAGATCCCGACAAC
TATACCCCCGCCAATCCTCTAAGCACCCCTCCCCATATCAAACCTGAATGGTACTTCCTATTTGCATATGCAATCCTCCGATCTAT
TCCCAATAAACTAGGAGGAGTTCTAGCCCTAGTTCTATCCATTTTAATCTTAGCAATTATCCCTGCCCTCCACACTTCCAAACAGC
GAGGAATAATGTTTCGACCACTAAGTCAATGCTTATTCTGATTCCTAGTAGCGGACCTTCTGACCCTGACATGAATTGGTGGCC
AACCTGTAGAACACCCCTTCATCACCATCGGCCAACTAGCCTCCATCCTATACTTCTCCATTCTTCTAATCCTAATACCCATCTCA
GGCATTATCGAAAACCGCCTCCTCAAA

>ZAM_T256; 16258-16327; +; trnT(aca)
GTCTTCGTAGTATATAGAATACCTTGGTCTTGTAAACCAAAAAAGGAGAACGCGTACCCTCCCTAAGACT

>ZAM_T256; 16328-16394; -; trnP(cca)
CAAGGAATAGTTTAAGAAAGAATTTCAGCTTTGGGTGCTGATGGTGGGGCTATTGCTTCTTCCTTGA
